# Nucleotide-mediated allosteric regulation of bifunctional Rel enzymes

**DOI:** 10.1101/670703

**Authors:** Hedvig Tamman, Katleen Van Nerom, Hiraku Takada, Niels Vandenberk, Daniel Scholl, Yury Polikanov, Johan Hofkens, Ariel Talavera, Vasili Hauryliuk, Jelle Hendrix, Abel Garcia-Pino

**Affiliations:** Cellular and Molecular Microbiology, Faculté des Sciences, Université Libre de Bruxelles (ULB), B-6041, Gosselies, Belgium; Department of Molecular Biology, Umeå University, SE-901 87 Umeå, Sweden; Laboratory for Molecular Infection Medicine Sweden (MIMS), Umeå University, SE-901 87 Umeå, Sweden; Molecular Imaging and Photonics, Chemistry Department, KU Leuven, Celestijnenlaan 200F (Chem&Tech), B-3001 Leuven, Belgium; Dynamic Bioimaging Lab, Advanced Optical Microscopy Centre and Biomedical Research Institute, Hasselt University, Agoralaan C (BIOMED), B-3590 Hasselt, Belgium; SFMB, Université Libre de Bruxelles (ULB) CP206/02, Boulevard du Triomphe, building BC, B-1050 Brussels, Belgium; Department of Biological Sciences, College of Liberal Arts and Sciences, University of Illinois at Chicago, 900 South Ashland Avenue, MBRB 4170, Chicago, IL 60607, USA; CMMI-Université Libre de Bruxelles, (ULB), B-6041, Gosselies, Belgium; WELBIO, Avenue Hippocrate 75, 1200 Brussels, Belgium

## Abstract

Bifunctional Rel stringent factors, the most broadly distributed class of RSHs, are ribosome-associated enzymes that transfer a pyrophosphate group from ATP onto the 3′ of GTP or GDP to synthesize (p)ppGpp and also catalyse the 3′ pyrophosphate hydrolysis of the alarmone to degrade it. The precise regulation of these enzymes seems to be a complex allosteric mechanism, and despite decades of research, it is unclear how the two opposing activities of Rel are controlled at the molecular level. Here we show that a stretch/recoil guanosine-switch mechanism controls the catalytic cycle of *T. thermophilus* Rel (Rel_*Tf*_). The binding of GDP/ATP stretches apart the NTD catalytic domains of Rel_*Tf*_ (Rel_*Tt*_^NTD^) activating the synthetase domain and allosterically blocking the hydrolase active site. Conversely, binding of ppGpp unlocks the hydrolase domain and triggers recoil of both NTDs, which partially buries the synthetase active site and precludes the binding of synthesis precursors. This allosteric mechanism acts as an activity switch preventing futile cycles of alarmone synthesis and degradation.

---

The cellular level of the bacterial alarmone (p)ppGpp^1,2^ – a key regulator of virulence and antibiotic tolerance – is controlled by the action of RelA/SpoT Homologue (RSH) enzymes^3-5^. The synthesis of (p)ppGpp involves transfer of the pyrophosphate group of ATP onto the 3′ of GDP or GTP, and these molecules are degraded by removal of the 3′ pyrophosphate moiety. The paradigm of RSH catalysis is based on the structure of *S. dysgalactiae* Rel (Rel_*Seq*_^NTD^) solved more than a decade ago. Rel_*Seq*_^NTD^ is formed by two catalytic domains with opposing activities – ppGpp hydrolase (HD) and ppGpp synthetase (SYN)^6^. Serendipitously, the two Rel_*Seq*_^NTD^ molecules observed in the same crystal lattice were locked in contrasting conformations leading to the hypothesis of reciprocal regulation of SYN and HD domains in archetypical RSH enzymes. However, the enzyme contained a nucleotide bound in each active site in one of the conformations and only one active site occupied in the other one. This contradicts earlier observations that suggested catalysis was incompatible with the simultaneous activation of synthetase and hydrolase function^7,8^. Thus, to directly test this hypothesis it is essential to solve the structures of Rel enzymes with catalytically engaged SYN or HD domains. To understand how nucleotide binding stimulates the enzymatic capacity of RSH enzymes, we took advantage of *T. thermophilus* Rel NTD (Rel_*Tt*_^NTD^, amino acid positions 1-355) as an experimental system. Rel_*Tt*_^NTD^ hydrolysis activity is virtually undetectable at 4°C (**Supplementary Fig. 1a** and **Supplementary Table 1**), which is not surprising given that *T. thermophilus* has an optimal growth temperature of about 65°C. This enabled co-crystallization in the presence of the native ppGpp substrate. To generate the structure of Rel_*Tt*_^NTD^ engaged in ppGpp synthesis, we used APPNP, a β-γ phosphate non-hydrolysable ATP analogue, that reacts exceedingly slowly with GDP in the active center of Rel_*Tt*_^NTD^ yielding ppGp p (**Supplementary Fig. 1b-c** and **Supplementary Table 1**).

The structure of unbound Rel_*Tt*_^NTD^ (**Fig. 1a** and **Supplementary Fig. 2a-d**) recapitulates the structures of Rel_*Seq*_^NTD^, *M. tuberculosis* Rel^9^ (RelA_*Mtb*_^NTD^) as well as the mono-functional synthetase-only *E. coli* RSH RelA (RelA_*Ec*_). Only the conformation of α-helices α6, α7 and loop α6-α7 are noticeably different (**Supplementary Fig. 2c-d**). In RelA_*Ec*_ the α6-α7 loop is projected towards the pseudo-hydrolase site of RelA_*Ec*_, effectively blocking the site, whereas in Rel_*Tt*_^NTD^, RelA_*Mtb*_^NTD^ and Rel_*Seq*_^NTD^ α6-α7 is partially disordered and pointing in an opposite direction^6,10-12^ (**Supplementary Fig. 2e-f**). Nucleotide-free Rel_*Tt*_^NTD^ contains two different conformations in the same crystal lattice with a similar arrangement of both catalytic domains. The major difference between both conformations is that the α6-α7 motif of the hydrolase domain is observed projected completely away from the active site with α-helix α6 largely unwound in an extended conformation (**Supplementary Fig. 2c-d** and **2f**). Although this extreme conformation is primarily stabilized by the lattice contacts and likely to be a result of the crystallization process, it underscores the highly dynamic nature of the hydrolase domain, in particular the α6-α7 motif and its importance for catalysis.

**Fig. 1.**
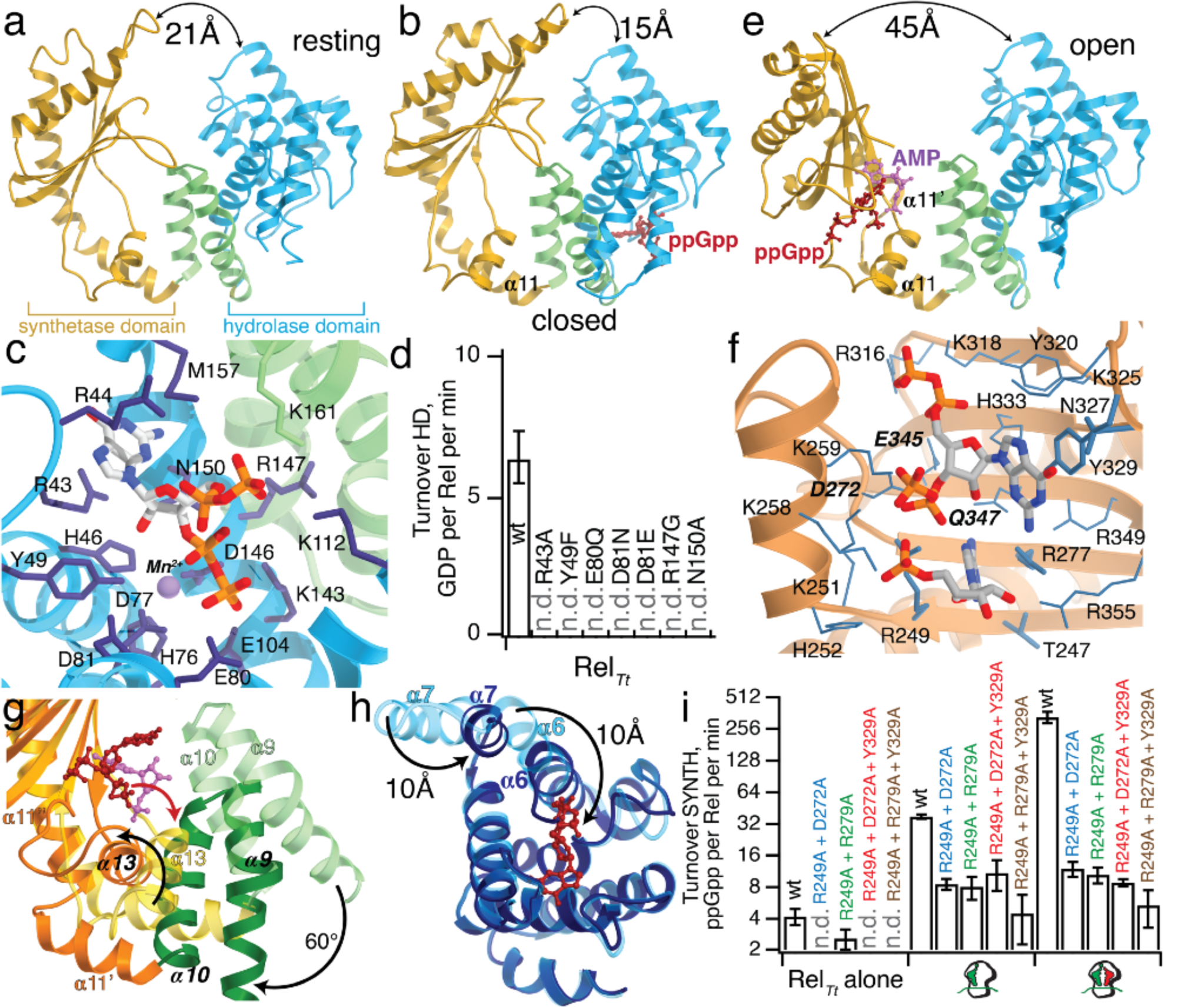
Structure of the different Rel_*Tt*_^NTD^ catalytic states. (**a**) Structure of Rel_*Tt*_^NTD^ in the resting (nucleotide free) state. The HD-domain is coloured in light blue, the α9-α10 α-helical substructure connecting the HD-and the SYN-domains is coloured in light green and the SYN-domain is coloured orange (**b**) Structure of Rel_*Tt*_^NTD^ in the active hydrolase state (close state, coloured as in panel (**a**)), bound to ppGpp (shown in red). (**c**) Details of the Rel_*Tt*_^NTD^-ppGpp binding interface, all important catalytic residues as well as the Mn2+ ion are labeled in the figure. (**d**) Impact of active site substitutions on the hydrolase activity of Rel_*Tt*_^NTD^, based on the interactions observed in the crystal structure of the Rel_*Tt*_^NTD^-ppGpp complex. In every case substitutions affecting the direct interaction with the guanosine base (R43A), the ribose (Y49F), residues involved in catalysis (E80Q, D81N, D81E and N150A) or interfering with the accommodation of the phosphate groups of (p)ppGpp (R147G) abrogate hydrolysis. (**e**) Structure of Rel_*Tt*_^NTD^ in the active synthetase state (open state, coloured as in panel (**a**)), bound to ppGpNp (shown in red) and AMP (shown in purple). (**f**) Details of the Rel_*Tt*_^NTD^-ppGpNp-AMP binding interface, all important catalytic residues are labeled in the figure with residues directly involved in catalysis shown in bold. (**g**) Superposition of the SYN-domain of the close (in light colors) and open (in dark colors) states illustrating the rigid body rotation movement of the α9-α10-α13 ‘transmission core’ induced by the binding of nucleotides, that results in the opening of the enzyme. (**h**) Superposition of the HD-domain of the close (light blue) and open (dark blue) states showing the allosteric changes triggered by the activation of the SYN-domain (HD partial occlusion) and the activation of the HD-domain (relocation of the α6-α7 snaplock). (**i**) ppGpp synthetase activity of Rel_*Tt*_, and R249A/R277A, R272A/R277A, R249A/R277A/Y329, R272A/R277A/Y329 substituted versions alone, and activated by either *T. thermophilus* 70S ribosome initiation complex (70S IC) or the 70S IC with deacylated tRNAVal in the A-site.

From the structure of the Rel_*Tt*_^NTD^-ppGpp complex (**Fig. 1b** and **Supplementary Fig. 3**), it is immediately apparent that the complex is in a more compacted conformation than the resting state of Rel_*Tt*_^NTD^ diverging from all other known conformation of Rel and RelA enzymes (**Fig. 1a** and **1b**). In the complex, ppGpp makes a large number of contacts with the enzyme (**Fig. 1c**) and is bound in a conformation reminiscent of that of ppG2′:3′p in the active site of Rel_*Seq*_^NTD^ and NADP bound to a single-domain small alarmone hydrolase (SAH) RSH hMesh1^13^ (PDBID 5VXA) (**Supplementary Fig. 4a-b**). The guanosine base is stacked between R43, R44 and M157 and makes hydrogen-bonds with S45, N150 and T153, while the ribose makes a van der Waals interaction with N150 and a hydrogen bond with Y49. The 2′- and 3′ oxygen atoms from the ribose are held very close (within 4.5 Å) to the Mn^2+^ ion and N150, which suggests an essential role for the metal ion in the deprotonation of the scissile bond and subsequent stabilization of a nucleophilic water molecule (**Fig. 1c**).

The hydrolase active site has a remarkable distribution of surface electrostatics. The site consists of a deep and wide cavity with one half of the site positively charged and involved in the stabilization of the 5′ poly-phosphate groups of the substrate and the other predominately acidic and more directly involved in the 3′-pyrophosphate hydrolysis (**Supplementary Fig. 4c**). The 3′-pyrophosphate group of ppGpp is bound in nearly the same position as that of ppG2′:3′p (**Supplementary Fig. 4a**) in the acidic half of the active site formed by D77, E80, D81, E104 and D146 which is crucial for catalysis. This acidic section of the active site has already been highlighted as a functional hot-spot^6^. Substitutions in the conserved HDX_3_ED motif^14^ (GD or EV) completely abrogate hydrolysis in Rel_*Seq*_^NTD6^ and the motif has diverged to _82_FPLADA_87_ in the inactive hydrolase domain of *E. coli* RelA suggesting this could be one of the factors contributing to the lack of activity of the specialised *E. coli* synthetase. Indeed, conservative substitutions of E80 (E80Q) and D81 (D81N) or even changes in the distance of the carboxylate group relative to ppGpp in the HD domain (as in D81E) have a strong effect on the activity of the enzyme (**Fig. 1d**).

The 5′-pyrophosphate group of ppGpp is stabilized by the damping effect of K112, K143, R147 and K161 and projects towards the α-helix α6 (**Fig. 1c**). From this binding mode we propose that the relative spatial arrangement between α6 and α9 would determine the specificity of the hydrolase function. Indeed, as observed in the Rel_*Seq*_^NTD^-ppG2′:3′p and hMesh1-NADP complexes, it is the local disposition of α6 and α9 what allows accommodation of the 5′-pyrophosphate and nicotinamide riboside groups (**Supplementary Fig 4a-b**). As with the case of the acidic patch, substitutions at this positive site (R147G), towards the phosphate binding region also strongly affect hydrolysis (**Fig 1d**).

The other crucial observation from the Rel_*Tt*_^NTD^-ppGpp complex is that the entire SYN domain moves 6 Å closer to the HD domain compared to the resting state, sterically occluding and switching off the synthetase active site (**Fig. 1a-b Supplementary Fig 5a-b**). In addition, the α6-α7 loop becomes fully structured and moves around 85° away from the position occupied by α6-α7 in RelA_*Ec*_ in the pseudo-active site of the HD domain. This movement of α6-α7 aligns together all the residues that constitute the aforementioned positive patch and allows the binding ppGpp in the active site. As a result, the hydrolase site of the enzyme becomes fully accessible in contrast with the synthetase site that is even more confined (**Supplementary Fig. 5a-b**).

To understand how substrates control the ppGpp synthesis by the SYN domain, we solved the structure of Rel_*Tt*_^NTD^ in a post-catalytic (PC) state (**Fig. 1e**) in complex with the reaction products AMP and ppGp_N_p (**Fig. 1f** and **Supplementary Fig. 6**). The structure of Rel_*Tt*_^NTD^ in this PC conformation (**Fig. 1e**) shows a remarkably contrasting picture with the Rel_*Tt*_^NTD^-ppGpp complex (**Fig. 1b**). The presence of two nucleotides in the synthetase site is accompanied by major rearrangements that stretch both domains almost 45 Å apart (**Fig. 1e**). The comparison of the two opposing conformations of the two active catalytic states suggest a potential allosteric signal transduction route (**Fig. 1g**). The central 3-α-helix bundle (C3HB) motif of the Rel_*Seq*_^NTD^ enzyme forms a small hydrophobic core with α13 of the SYN domain that connects both catalytic domains^6^. The wedging effect of the nucleotides is thus spread towards the HD domain via the α13-C3HB ‘transmission’ core that has swivelled orthogonally to the α9 dipole approximately 60° (**Fig. 1g** and **Supplementary Fig. 6b-c**). This fractures α11 in two (α11’ and α11’’), exposing the SYN domain active site and stretching the HD domain away. This conformational rearrangement portrays a much bigger allosteric effect associated with the switch to an active synthetase state than that expected from the two conformations of Rel_*Seq*_^NTD^ which involved a rotation of 10° stretching both domains around 2.4 Å apart^6^. Considering the lattice constraints and the partial occupancy of nucleotides in both active sites of Rel_*Seq*_^NTD^, it is not surprising that the enzyme is observed in an intermediate state closer to the resting state than to either active catalytic conformation.

In the HD domain, these rearrangements are accompanied by a conformational change in loop α6-α7 that approaches towards the catalytic residues of the hydrolase centre (**Fig. 1h**). These structural changes effectively close the hydrolase site, reducing its radius to almost half its size and expanding the radius and dimensions of synthetase active site in order to accommodate both nucleotide substrates (**Supplementary Fig. 6b-c**). From the structure of the free Rel_*Tt*_^NTD^ and Rel_*Seq*_^NTD^ bound to GDP, it is clear that only the presence of both nucleotides in the active site of the synthetase domain can trigger such large domain rearrangements.

The PC active site of Rel_*Tt*_^NTD^ resembles the pre-catalytic state observed in the structure of the RelP SAS enzyme from *S. aureus*, a single-domain (p)ppGpp synthetase-only RSH enzyme that lacks additional catalytic or regulatory domains. The overall interactions of ppGp_N_p with the synthetase active site are similar to those observed in the RelP-GTP-APCPP (PDBID 6EWZ) and RelP-pppGpp (PDBID 6EX0) complexes^15^ (**Fig. 1e, Supplementary Fig. 7a-b**). Only small differences in the orientation of the G-loop^16^ and the accommodation of the ligand are observed due to the lack of the additional phosphate group that is present in both RelP complexes (**Supplementary Fig. 7b)**. The coordination of the adenosine group of AMP in the active site also resembles that of the pre-catalytic RelP with the adenosine base stacked between R249 and R277 and the α-phosphate of AMP coordinated in the same manner as that observed in the RelP-GTP-APCPP complex by R249 and K215 (**Supplementary Fig. 7b)**. In addition, the β1-α13 loop and α13 contribute a patch of positive residues that stabilize the pyrophosphate group transferred to ppGp_N_p, which is around 2.0 Å away from the site it would occupy in a pre-catalytic state, similar to that of RelP, as part of APCPP. These two active site elements are observed in a similar orientation to that of the β1-α2 loop and α2 of RelP, coordinating the tri-phosphate groups of APCPP (**Supplementary Fig 7b**). Indeed mutations that destabilize ATP binding and affect the interaction of GDP/GTP with the G-loop (R249A, D272, R277A and Y329A) have a strong impact on the activity of the native full length Rel_*Tt*_ when it is assayed by itself or when the enzyme is activated by either *T. thermophilus* 70S ribosome initiation complex (70S IC) or the 70S IC with deacylated tRNA^Val^ in the A-site, the ultimate natural activator of ppGpp synthesis by Rel_*Tt*_ (**Fig 1i**).

Our structural data suggest that the presence of nucleotides in either the hydrolase or the synthetase domain would prime the enzyme for that particular function, switching off the other catalytic site. Such an allosteric effect would manifest itself in the form of changes in the width of the conformational landscape that would tilt the dynamic equilibrium of the population ensemble towards the favoured state as a function of the concentration of the nucleotides in solution. We directly challenge this hypothesis using single-molecule fluorescence resonance energy transfer (smFRET). For this, we constructed Rel_*Tt*_^NTD^_6/287_, a variant of Rel_*Tt*_^NTD^ that allows fluorescent labels to be attached at cysteine residues introduced at positions 6 and 287 (**Fig. 2a**). The FRET-averaged inter-dye distance ⟨R_DA_⟩_E_ predicted for Rel_*Tt*_^NTD^_6/287_ based on our crystal structures is 75 Å for the open form (Rel_*Tt*_^NTD^-AMP-ppGp_N_p complex) and 57 Å for the closed form (Rel_*Tt*_^NTD^-ppGp_N_p complex).

**Fig. 2.**
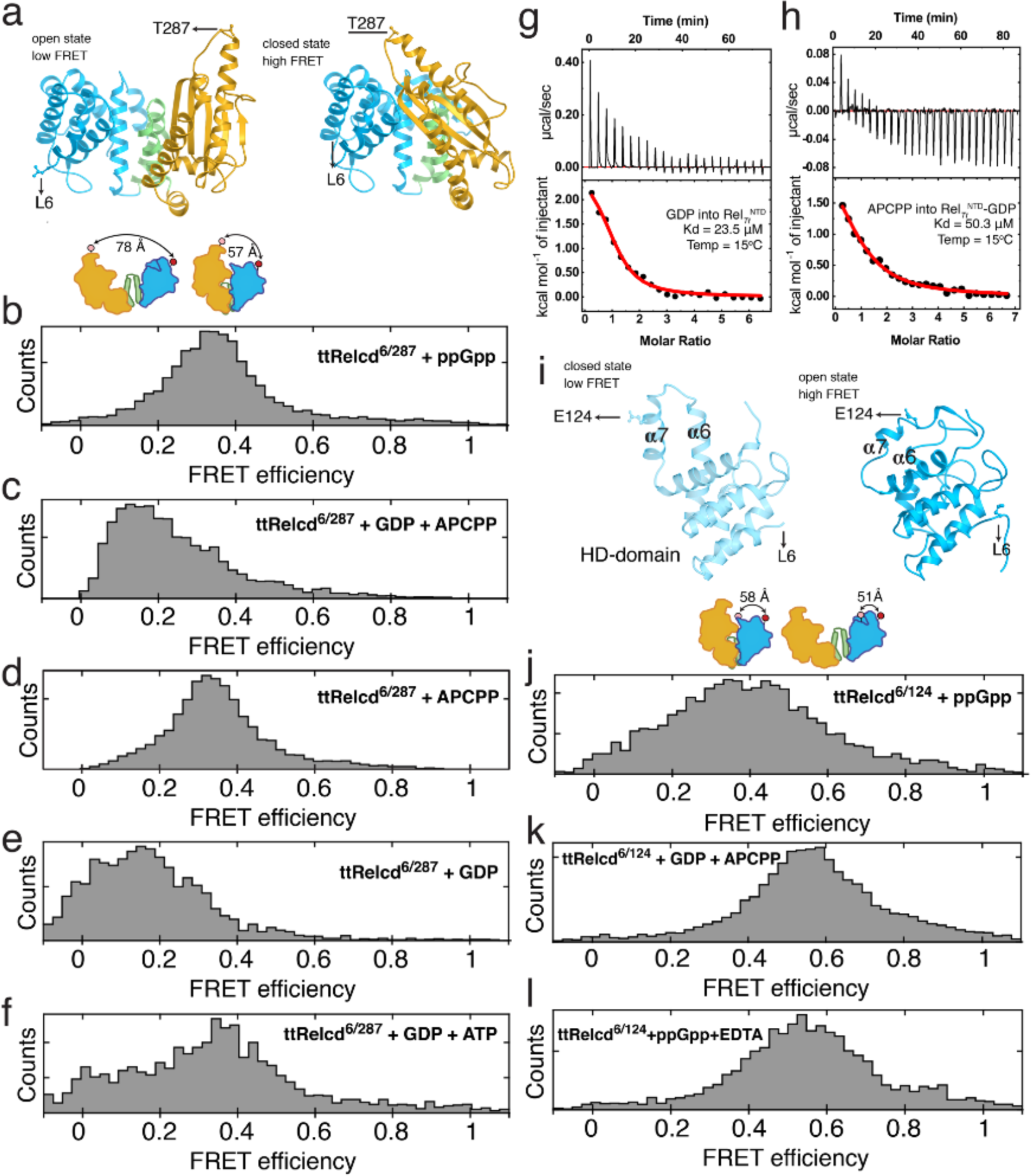
Rel_*Tt*_^NTD^ conformational dynamics in the presence of nucleotides assessed by smFRET. **(a)** Structural models of Rel_*Tt*_^NTD^ in the open and closed conformations (to probe the inter-domain movements associated with the nucleotides allosteric control). Dye attachment sites are Leu6Cys and Thr287Cys. 1D projection histograms of the FRET efficiency of the Rel_*Tt*_^NTD^_6/287_ in the presence of ppGpp (**b**), GDP+APCPP (**c**), APCPP (**d**), GDP (**e**) and GDP+ATP (**f**). The analysis of the smFRET data suggest that the high FRET state of the enzyme in the presence of ppGpp and APCPP is consistent with the closed form observed in the crystal structure of the complex with ppGpp. By contrast the low FRET state of the enzyme observed in the presence of GDP and GDP+APCPP is consistent with the open state of the enzyme observed in the structure of Rel_*Tt*_^NTD^-ppGp_N_p-AMP. (**g**) Titration of GDP into Rel_*Tt*_^NTD^. (**h**) Titration of APCPP into Rel_*Tt*_^NTD^ in the presence of saturating amounts of GDP. (**i**) Structural models of Rel_*Tt*_^NTD^ in the open and closed conformations. Dye attachment sites are Leu6Cys and Thr124Cys (to probe HD-domain movements associated with the nucleotides allosteric control). 1D projection histograms of the FRET efficiency of the Rel_*Tt*_^NTD^_6/124_ in the presence of ppGpp (**j**), GDP+APCPP (**k**) and ppGpp+EDTA (**l**). The lower FRET state of the hydrolase domain in the presence of ppGpp is consistent with the movement of α-helices α6 and α7 which allow the binding of the substrate into the active site. By contrast the higher FRET state observed in the presence of GDP+APCPP is compatible with the snaplock switch that closes the hydrolase state coupled to the activation of the synthetase domain and the overall stretching of both catalytic domains.

While in the presence of ppGpp, Rel_*Tt*_^NTD^_6/287_ shows a homogenous population with ⟨R_DA_⟩_E_ of 64 Å (**Fig. 2b** and **Supplementary Fig 8a**), when incubated with GDP combined with the non-hydrolysable ATP analogue – APCPP – the Rel_*Tt*_^NTD^_6/287_ population shifted to a ⟨R_DA_⟩_E_ of 72 Å consistent with the opening of the enzyme (**Fig. 2c** and **Supplementary Fig 8b**). These clearly different conformational states matched the theoretical states expected from the aforementioned crystal structures (**Fig. 1b** and **Fig. 1e**). In the presence of APCPP alone, Rel_*Tt*_^NTD^_6/287_ remained in the closed conformation (**Fig. 2d** and **Supplementary Fig 8c**). Conversely, GDP triggered the opening of the enzyme (⟨R_DA_⟩_E_ of 70 Å) (**Fig. 2e** and **Supplementary Fig 8d**). Interestingly, when ATP was added after pre-incubating the enzyme with GDP, the ppGpp produced as a result of the reaction returned the equilibrium to the closed state (**Fig. 2f** and **Supplementary Fig 8e**). This suggests the reaction occurs in a sequential manner with the guanosine substrates binding first in the active site.

To directly address the sequential binding hypothesis, we monitored the binding of these nucleotides using Isothermal Titration Calorimetry (ITC). GDP binds Rel_*Tt*_^NTD^ with an affinity of 23 μM (**Fig. 2g** and **Supplementary Table 2**) and in excellent agreement with the smFRET data APCPP alone does not bind Rel_*Tt*_^NTD^ (**Supplementary Fig. 9a** and **Supplementary Table 2**). However, once the Rel_*Tt*_^NTD^-GDP complex is formed, APCPP binds with an affinity of around 50 μM (**Fig. 2h** and **Supplementary Table 2**). The incorporation of both GDP and APCPP is entropically driven (**Supplementary Fig 9b** and **Supplementary Table 2**). In the case of the binding of GDP to the SYN-domain, the release of ordered water molecules from the GDP binding cleft is accompanied by the movement of α6-α7 that partially occludes the HD active center and an increase in the enzyme flexibility as observed in the crystal structure. All these structural changes are consistent with entropically driven binding events. The smFRET data shows that GDP binding is also coupled to the opening of the enzyme which reveals the otherwise buried ATP biding site. In addition, it creates an ‘entropy reservoir’ that drives the binding of APCPP with the the release of water molecules from the now exposed binding cleft. This supports the notion that GDP must ‘open’ the enzyme to reveal the ATP binding site, as predicted from the analysis of the crystallographic data.

We hypothesized that the motions of the α6-α7 loop were coupled to the allosteric switching of the enzyme, constituting a crucial element of the intramolecular crosstalk between domains. In the hydrolase-ON (Rel_*Tt*_^NTD^-ppGpp complex, close state), this loop was projected away from the hydrolase active site allowing the binding of ppGpp whereas in the synthetase-ON state (Rel_*Tt*_^NTD^ post-catalytic open state) α6-α7 moved towards the hydrolase active site precluding the binding of ppGpp, effectively switching off the hydrolase function. We used Rel _*Tt*_^NTD^ _6/124_, a Rel_*Tt*_^NTD^ variant fluorescently labelled via cysteine residues introduced at residue positions 6 and 124 (**Fig. 2i**). In the presence of ppGpp, Rel_*Tt*_^NTD^ _6/124_ is observed in a low FRET state of ⟨R_DA_⟩_E_ 62 Å, indicating displacement of the loop away from the active site (**Fig. 2j** and **Supplementary Fig. 8f**). Conversely, when bound to GDP and APCPP, the enzyme switched to a high FRET state (⟨R_DA_⟩_E_ of 55 Å) indicative of the loop movement towards the active site (**Fig. 2k** and **Supplementary Fig. 8g**). These smFRET data are in good agreement with ⟨R_DA_⟩_E_ estimates based on the structural data that predicts a distance between dyes of 51 Å for the open state and 60 Å for the closed. The removal of the Mn^2+^ ion from the hydrolase site by incubation with EDTA, precluded the close conformation even in the presence of ppGpp (**Fig. 2l** and **Supplementary Fig 8h**). These observations support the role of α6-α7 as an allosteric ‘snaplock’. When the snaplock is stabilized away from the hydrolase active site, the enzyme is predominantly in the closed state and in the hydrolase-ON conformation. By contrast, releasing the snaplock leads to closure of the hydrolase active site and the ‘snap’-opening of the enzyme exposing the synthetase active site (**Supplementary Fig 6b-c**).

The dynamic modulation of the cellular alarmone levels is paramount to maintenance of cellular homeostasis. Here we showed that Rel enzymes possessing active hydrolase and synthetase domains rely on additional levels of allosteric regulation besides the control that the C-terminal regulatory domains exert, which prevent the occurrence of futile catalytic cycles. The activation of one of the catalytic domains entails the physical blockade and active site misalignment of the other. Our results support the view that the allosteric motion of α-helices α6 and α7 is coupled to the catalytic cycle precluding or allowing the access of substrate to the hydrolase active site whereas the relative conformational state between domains regulates the synthetase function, hindering access to the synthetase site in the close state. This allosteric control provides a *bona fide* on/off switch that renders one domain completely blocked while the other is active (**Fig. 3**). In addition, our results suggest that the allosteric control of long RSH enzymes may differ from that of SASs such RelP or RelQ that also bind their substrates in an ordered sequence but incorporate first with ATP and then GDP or GTP^17^. We show that in long RSH enzymes this sequence is reversed. This divergence in the catalytic mechanism is most likely due to the loss of the hydrolase domain and with it the requirement of triggering an open state in SASs to bind ATP.

**Fig. 3.**
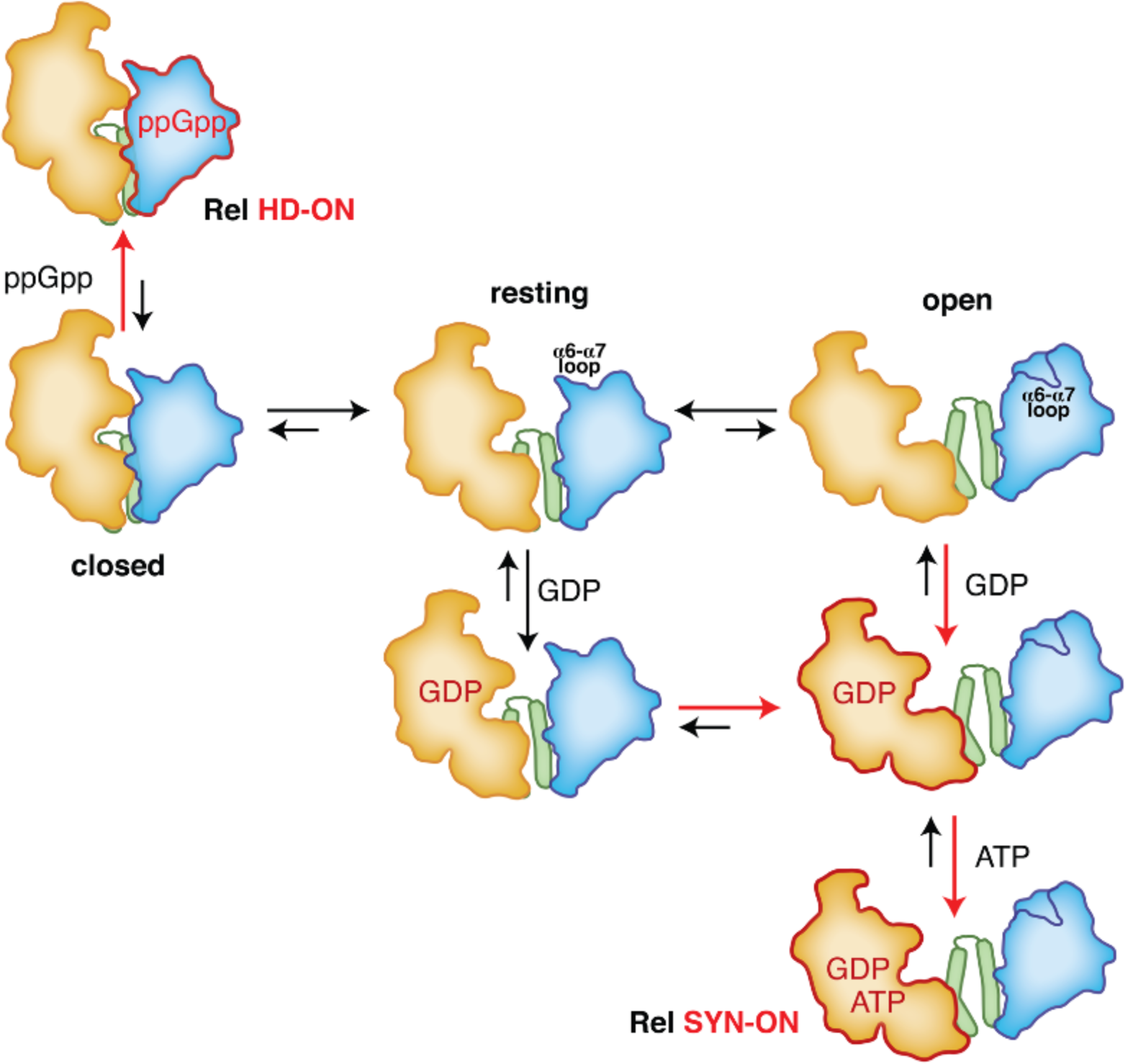
Cartoon representation of the molecular model for the mechanism of regulation of Rel_*Tt*_^NTD^ catalysis as a function of nucleotides. In absence of ligands the resting state of the enzyme is consistent with the conformations observed in RelA_*Mtb*_^NTD^ and RelA_*Ec*_ with the catalytic site of the SYN-domain is partially occluded (the ATP binding site completely buried in the resting state). The ppGpp synthesis cycle is initiated by GDP binding which triggers an open form that allows the consequent binding of ATP, followed by the synthesis of ppGpp. Moreover, the stabilization of this open state is coupled to changes in the active site of the HD-domain, that preclude hydrolysis. Conversely, ppGpp binding to the HD-domain triggers the overall closing of the enzyme resulting in the complete occlusion of the active site of the SYN-domain safeguarding against non-productive ppGpp synthesis and the alignment of the active site residues in the HD-domain to accommodate ppGpp and catalyze the hydrolysis of the 3′-pyrophosphate group of the molecule.

The current view of the role of nucleotides in the control of Rel enzymes, based on known structures, does not provide a conclusive explanation as to how the conformational interplay between the two catalytic domains is linked to the activation of one site to the detriment of the other. Moreover in all known structures of different RSH enzymes the two catalytic domains are observed in similar conformations that are incompatible with an active synthetase state^6,9-12^.

The regulation of catalysis of bifunctional enzymes is usually dominated by allosteric transitions preventing the occurrence of futile cycles^18^. The stretch/recoil mechanism we propose provides the basis for the macromolecular control of the enzyme by its opposing substrates. The synchronous action of this additional regulatory layer, together with the spatial control of the ribosome as the docking platform of the enzyme for (p)ppGpp synthesis^10-12^ or the PTS for ppGpp hydrolysis^19^, denote a tightly regulated mechanism that prevents the occurrence of futile catalytic cycles and facilitates the action of the alarmone as a bacterial phenotypic switch.

## Acknowledgements

We acknowledge the use of the synchrotron-radiation facility at the Soleil synchrotron Gif-sur-Yvette, France, under proposals 20150717, 20160750 and 20170756; we also thank the staff from Swing, PROXIMA-1 and PROXIMA-2A beamlines at Soleil for assistance with data collection. Prof. J. Hofkens is kindly thanked for the use the single molecule FRET facility at KULeuven. This work was supported by grants from the Fonds National de Recherche Scientifique, FNRS-EQP U.N043.17F, FRFS-WELBIO CR-2017S-03, FNRS-PDR T.0066.18 and the Joint Programming Initiative on Antimicrobial Resistance (JPI-EC-AMR -R.8004.18-) to A.G.-P. The Program ‘Actions de Recherche Concertée’ 2016–2021 and Fonds d’Encouragement à la Recherche (FER) of ULB, Fonds Jean Brachet and the Fondation Van Buuren to A.G.-P.; the Molecular Infection Medicine Sweden (MIMS), Swedish Research council (grant 2017-03783 to and Ragnar Söderberg foundation fellowship to V.H; J.Hendrix and J.Hofkens are grateful to the Research Foundation Flanders (FWO Vlaanderen, G0B4915N) and large infrastructure grant ZW15_09 GOH6316N) and the KU Leuven Research Fund (C14/16/053); J.Hofkens thanks financial support of the Flemish government through long term structural funding Methusalem (CASAS2, Meth/15/04). K.V.N. was supported by a PhD grant from the Fonds National de Recherche Scientifique FNRS-FRIA; N.V. acknowledges the Agency for Innovation by Science and Technology in Flanders (IWT) for a PhD grant. H.Tamman was supported by a Chargé de Recherches fellowship from the FNRS (CR/DM-392) and H.Takada was supported by the postdoctoral grant from the Umea □Centre for Microbial Research (UCMR).

## Author contributions

H.Tamman, K.V., N.V. D.S and A.T. performed biophysical, structural biology and smFRET experiments, H.Takada performed the biochemical assays on Rel_*Tt*_, YP was involved in the initial steps of the preparation of *T. thermophilus* ribosomes, J.Hendrix and J.Hofkens supervised the smFRET data analysis and V.H, J.Hendrix. and A.G.-P. designed and supervised research and wrote the paper.

## References

1. Cashel, M. & Gallant, J. Two compounds implicated in the function of the RC gene of Escherichia coli. Nature 221, 838–41 (1969).

2. Hauryliuk, V., Atkinson, G.C., Murakami, K.S., Tenson, T. & Gerdes, K. Recent functional insights into the role of (p)ppGpp in bacterial physiology. Nat Rev Microbiol 13, 298–309 (2015).

3. Atkinson, G.C., Tenson, T. & Hauryliuk, V. The RelA/SpoT homolog (RSH) superfamily: distribution and functional evolution of ppGpp synthetases and hydrolases across the tree of life. PLoS One 6, e23479 (2011).

4. Laffler, T. & Gallant, J.A. Stringent control of protein synthesis in E. coli. Cell 3, 47–9 (1974).

5. Stent, G.S. & Brenner, S. A genetic locus for the regulation of ribonucleic acid synthesis. Proc Natl Acad Sci U S A 47, 2005–14 (1961).

6. Hogg, T., Mechold, U., Malke, H., Cashel, M. & Hilgenfeld, R. Conformational antagonism between opposing active sites in a bifunctional RelA/SpoT homolog modulates (p)ppGpp metabolism during the stringent response [corrected]. Cell 117, 57–68 (2004).

7. Avarbock, D., Salem, J., Li, L.S., Wang, Z.M. & Rubin, H. Cloning and characterization of a bifunctional RelA/SpoT homologue from Mycobacterium tuberculosis. Gene 233, 261–9 (1999).

8. Mechold, U., Potrykus, K., Murphy, H., Murakami, K.S. & Cashel, M. Differential regulation by ppGpp versus pppGpp in Escherichia coli. Nucleic Acids Res 41, 6175–89 (2013).

9. Singal, B. et al. Crystallographic and solution structure of the N-terminal domain of the Rel protein from Mycobacterium tuberculosis. FEBS Lett 591, 2323–2337 (2017).

10. Arenz, S. et al. The stringent factor RelA adopts an open conformation on the ribosome to stimulate ppGpp synthesis. Nucleic Acids Res (2016).

11. Brown, A., Fernandez, I.S., Gordiyenko, Y. & Ramakrishnan, V. Ribosome-dependent activation of stringent control. Nature 534, 277–80 (2016).

12. Loveland, A.B. et al. Ribosome*RelA structures reveal the mechanism of stringent response activation. Elife 5(2016).

13. Sun, D. et al. A metazoan ortholog of SpoT hydrolyzes ppGpp and functions in starvation responses. Nat Struct Mol Biol 17, 1188–94 (2010).

14. Aravind, L. & Koonin, E.V. The HD domain defines a new superfamily of metaldependent phosphohydrolases. Trends Biochem Sci 23, 469–72 (1998).

15. Manav, M.C. et al. Structural basis for (p)ppGpp synthesis by the Staphylococcus aureus small alarmone synthetase RelP. J Biol Chem 293, 3254–3264 (2018).

16. Steinchen, W. et al. Structural and mechanistic divergence of the small (p)ppGpp synthetases RelP and RelQ. Sci Rep 8, 2195 (2018).

17. Steinchen, W. et al. Catalytic mechanism and allosteric regulation of an oligomeric (p)ppGpp synthetase by an alarmone. Proc Natl Acad Sci U S A 112, 13348–53 (2015).

18. Okar, D.A. et al. PFK-2/FBPase-2: maker and breaker of the essential biofactor fructose-2,6-bisphosphate. Trends Biochem Sci 26, 30–5 (2001).

19. Ronneau, S. et al. Regulation of (p)ppGpp hydrolysis by a conserved archetypal regulatory domain. Nucleic Acids Res 47, 843–854 (2019).

