## Supplementary Figures for "Nucleotide-mediated allosteric regulation of bifunctional Rel enzymes"

**Supplementary Fig. 1. Biochemical characterization of full-length Rel*Tt*.** (**a**) Hydrolysis activity of 100 nM full-length Rel*Tt* was assayed in the presence of 0.5 mM 3H-labelled ppGpp substrate either at 4˚C or 40˚C. Synthetic activity of 30 nM full-length Rel*Tt* was assayed at 40˚C in the presence of either 1 mM ATP (**b**) or 1 mM APPNP (**c**), as well 0.3 mM of 3H-labelled GDP and 0.1 mM ppGpp. As indicated on the figure, the reactions were supplemented with 120 nM *T. thermophilus* 70S IC(MV) added either alone or in combination with 2 µM deacylated *E. coli* $tRNA^{Val}$. All experiments were performed in HEPES:Polymix buffer, pH 7.5, 5 mM $Mg^{2+}$, either in the presence (**a**) or absence (**b** and **c**) of 0.5 mM $Mn^{2+}$. Error bars represent SDs of the turnover estimates by linear regression. All the reaction parameters are shown in **Supplementary Table 1**.

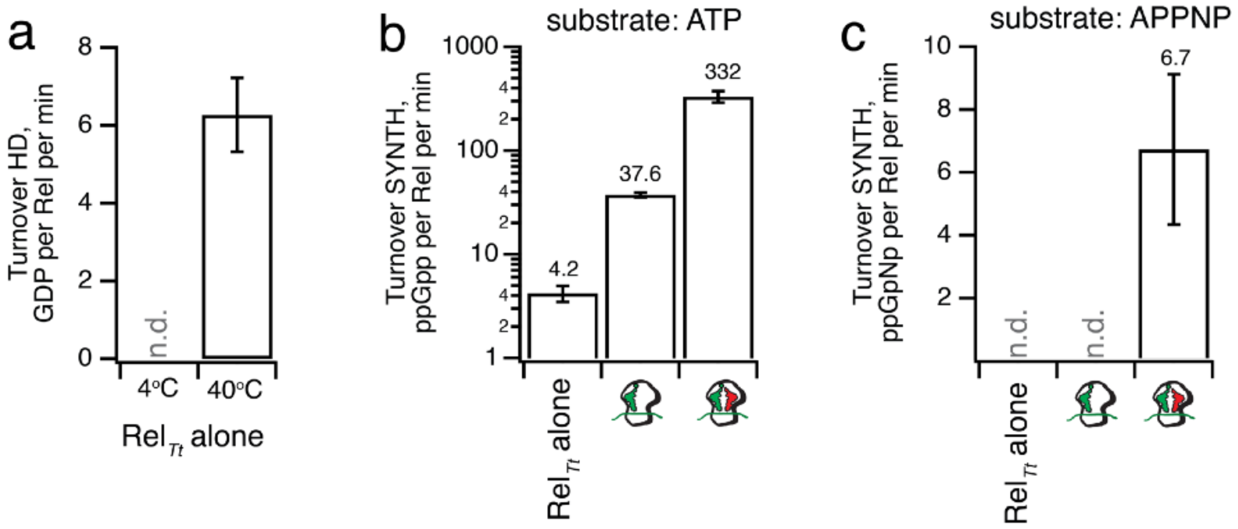

**Supplementary Fig. 2.** Conformations observed in the crystal structure of the unbound $\mathrm{Rel}_{Tt}^{\mathrm{NTD}}$ (resting state of the enzyme). (**a**) Lattice packing of the $P4_12_12$ crystals of free $\mathrm{Rel}_{Tt}^{\mathrm{NTD}}$. (**b**) Representation of the $\mathrm{Rel}_{Tt}^{\mathrm{NTD}}$ topology with the HD-domain coloured in light blue, the α9-α10 linker region in light green and the SYN-domain in light orange. (**c**) Conformation 1 (colored based on the topology panel), consistent with most of the conformations observed in the catalytic domains of Rel enzymes. (**d**) In conformation 2 (as in (**a**) but using dark shades) both catalytic domains are arranged as in conformation 1, however the α6-α7 motif (labeled in the figure) protrudes away from the structure and is stabilised by lattice contacts. (**d**) Superposition of the catalytic domains of different Rel-like enzymes in the resting state (from *T. thermophilus* (in green), *M. tuberculosis* (in yellow) and *S. dysgalactiae* (in purple). From the superposition it becomes apparent that the α6-α7 α-helical motif of the hydrolase domain accounts for the main differences in conformation between these structures. (**f**) Detailed view of the α6-α7 α-helical motif as in (**b**) but now including α6-α7 from Mesh1 (in red) a constitutively active ppGpp hydrolase from *H. sapiens*. (**g**) The alternative conformation observed in the lattice, shown in panel (**c**), is most likely the result of the lattice constrains on the fold. Nevertheless it underscores the dynamic nature of this catalytic domain and, this dynamic interplay involving the ppGpp binding site in the HD-domain must have a strong impact in catalysis.

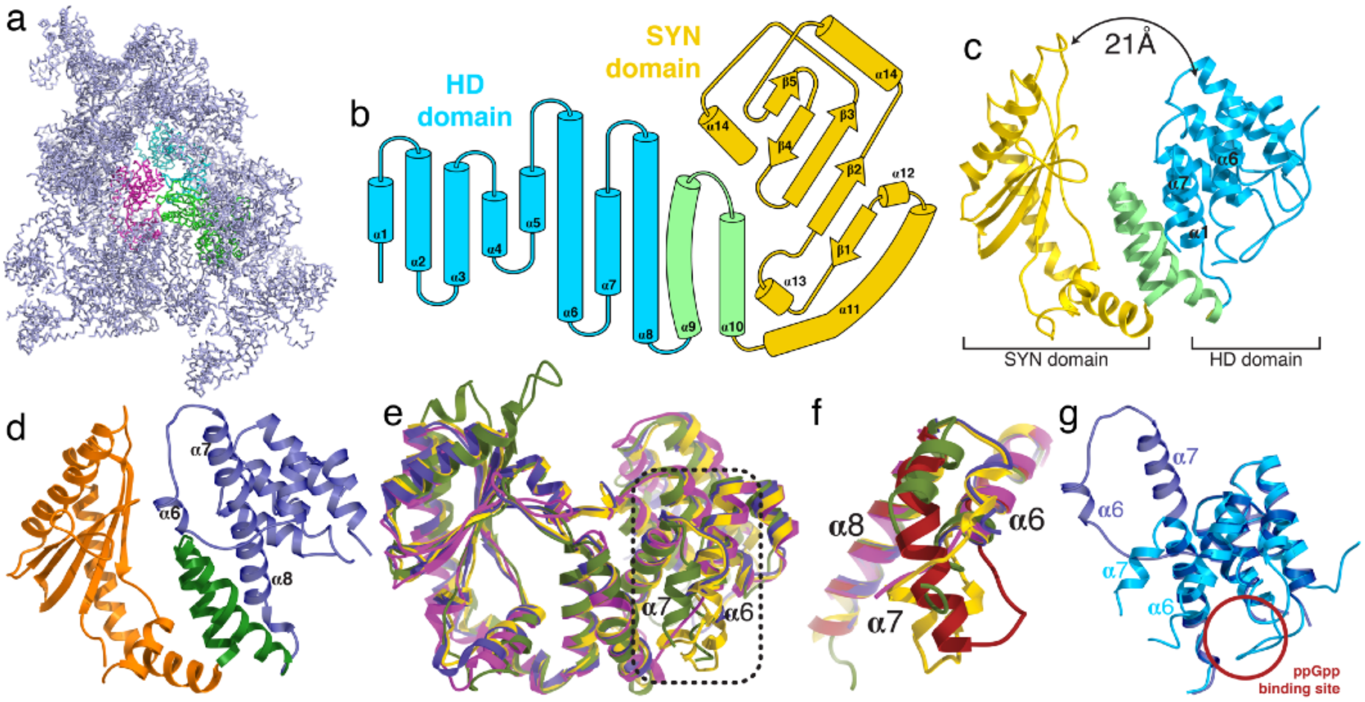

**Supplementary Fig. 3.** Electron density map representation (unbiased mFo-DFc), of the hydrolase domain Rel$_{Tt}^{NTD}$ as observed in the closed form (coloured in blue) bound to ppGpp (in atom colours), after refinement with Buster/TNT. The map was calculated from the MR solution omitting the ligand.

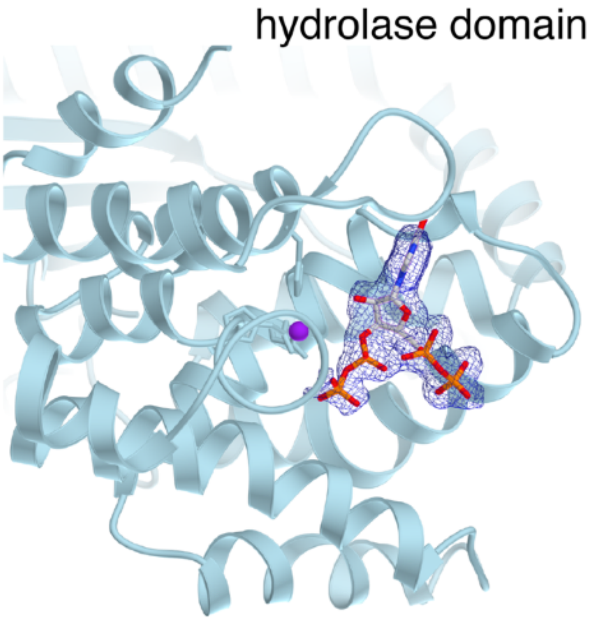

**Supplementary Fig. 4.** (**a**) Structure of the Rel$_{Tt}$$^{NTD}$-ppGpp complex (shown in light blue) superimposed on the Rel$_{Seq}$$^{NTD}$-ppG2':3'p complex, shown in red (**b**). Structure of the Rel$_{Tt}$$^{NTD}$-ppGpp complex (shown in light blue) superimposed on the Mesh1-NADP complex (shown in red). The comparison reveals a conserved active site architecture held together by the presence of a $Mn^{2+}$ ion that coordinates residues from three different α-helices (α3, α4 and α8) and is directly involved in catalysis. In addition these superpositions suggest a crucial role for α-helices α6 and α7 in the accommodation and stabilisation of the substrate in the active site. (**c**) Surface representation of Rel$_{Tt}$$^{NTD}$-ppGpp (shown in yellow). The surface is colored on the basis of electrostatic potential. The nucleotide binding site (for ppGpp or pppGpp) is traced in black dashed lines, highlighting the peculiar surface electrostatics of the active site with one acid half involved directly in hydrolysis and positive half involved in the stabilisation of the large number of phosphate groups carried by (p)ppGpp.

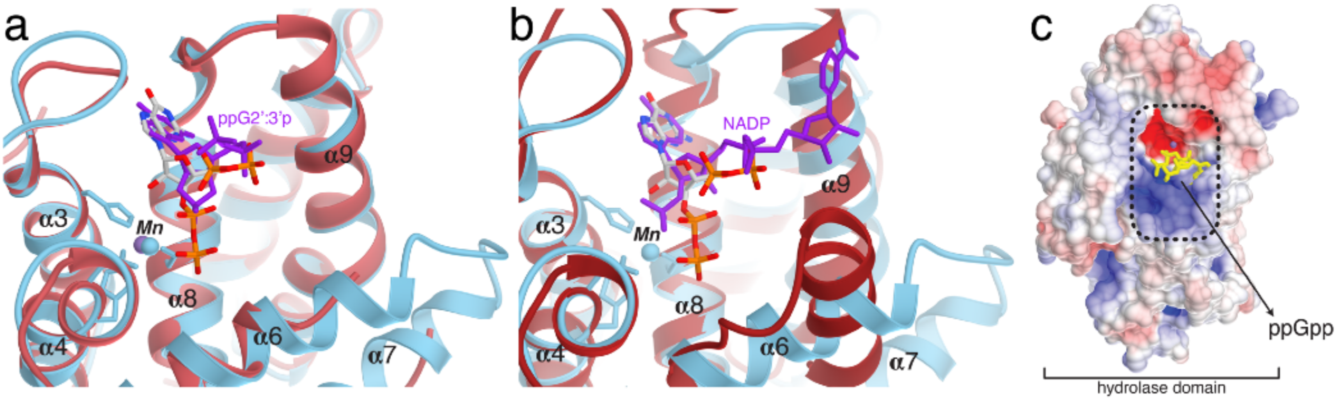

**Supplementary Fig. 5.** Allosteric rearrangements and active site reshaping of Rel$_{Tt}^{\mathrm{NTD}}$ as a function of nucleotides (colored as in **Fig. 1b**). (**a**) The hydrolase active site in the catalytically compatible form observed in the Rel$_{Tt}^{\mathrm{NTD}}$-ppGpp complex forms an L-shape crevice to accommodate (p)ppGpp (shown as a red volume). The allosteric arrangements involved in the active site setup are couple to the closing of the enzyme that constricts the synthetase active site (shown as a light blue volume) and prevents ppGpp synthesis. (**b**) Analysis of the hydrolase (solid symbols) and synthetase active site (open symbols) dimensions from the structures complexes of Rel$_{Tt}^{\mathrm{NTD}}$-ppGpp and Rel$_{Tt}^{\mathrm{NTD}}$-ppGpNp-AMP. In the Rel$_{Tt}^{\mathrm{NTD}}$-ppGpp complex the HD active site is larger and 2Å broader on average than when the enzyme is in the active synthetase form in contrast with what is observed in the synthetase site which becomes much larger and significantly broader in the Rel$_{Tt}^{\mathrm{NTD}}$-ppGpNp-AMP compared with the closed form. (**c**) Active site representation of the enzyme in the open form (Rel$_{Tt}^{\mathrm{NTD}}$-ppGpNp-AMP complex) coloured as in (**a**). The figure shows the stretching of the two catalytic domains involved in the correct arrangement of the synthetase catalytic site which is coupled to the closing and inactivation of the hydrolase domain catalytic site.

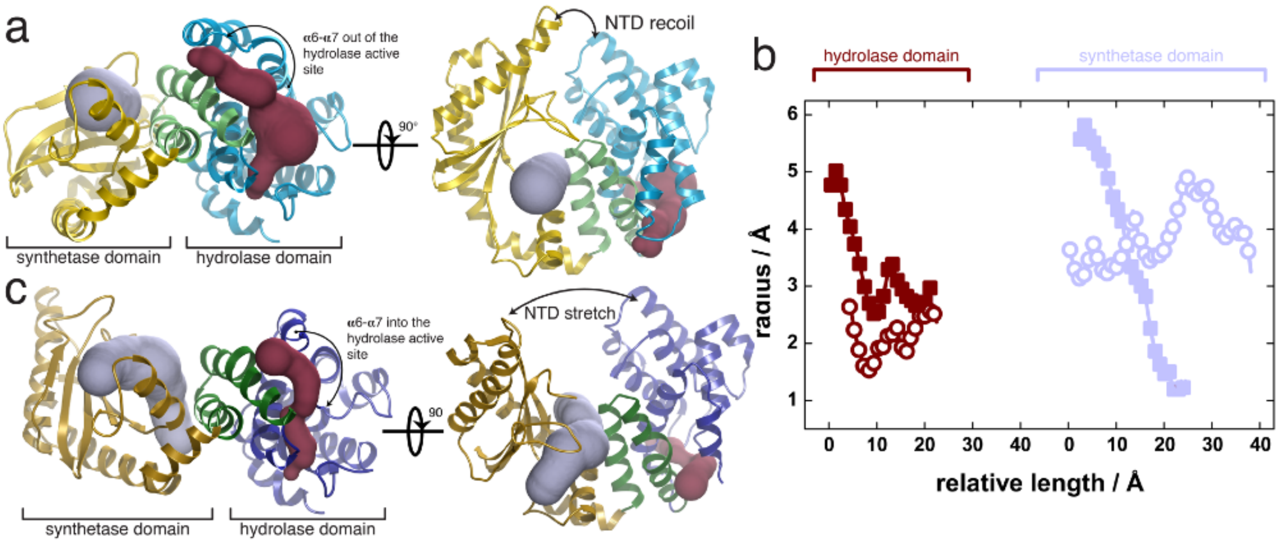

**Supplementary Fig. 6. (a)** Electron density map representation (unbiased mFo-DFc), of the synthetase domain $Rel_{Tt}^{NTD}$ as observed in the open form (coloured in yellow) bound to $ppGp_Np$ and AMP (in red and purple respectively), after refinement with Buster/TNT. The map was calculated from the MR solution omitting the ligand. **(b)** Superposition of the open (in dark colors) and closed conformations (in light colors) of $Rel_{Tt}^{NTD}$. The α9-α10-α13 'transmission core' is shown in magenta for the closed state and in green for the open state. The rigid body swivel of the transmission core triggers the opening of the enzyme and partial occlusion of the HD-domain active site. **(c)** Details of the conformational rearrangement of the transmission core induced by the binding of nucleotides.

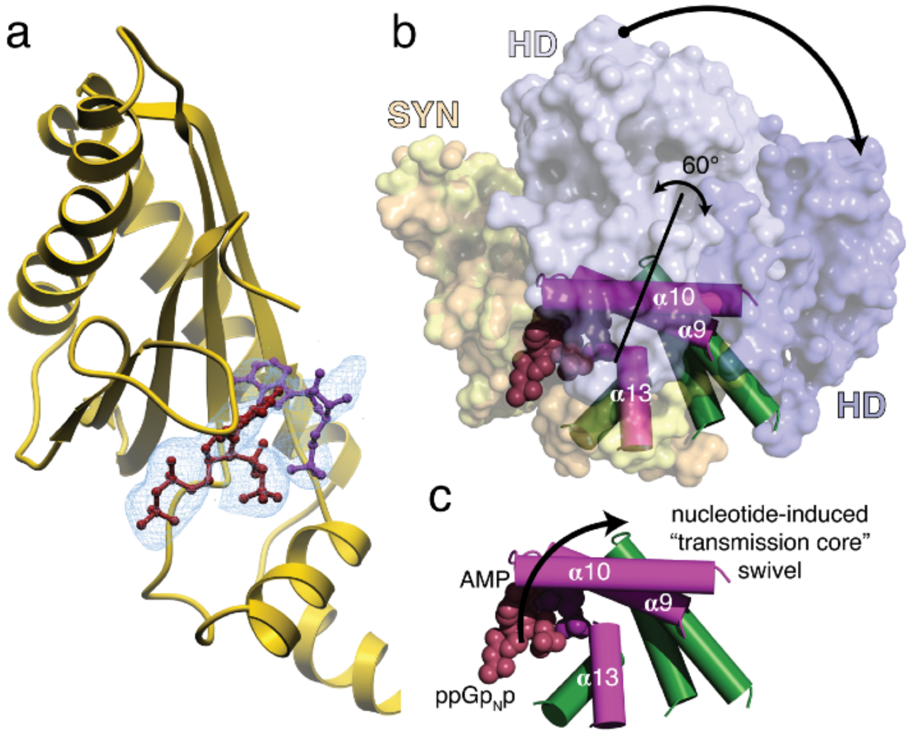

**Supplementary Fig. 7.** (**a**) Superposition of the active site residues of Rel$_{Tt}^{NTD}$ (in yellow) in the PC state on the active site residues of the RelP (in green) small alarmone syntetase in a pre-catalytic state (bound to GTP and APCPP). The catalytic residues of Rel$_{Tt}^{NTD}$ are shown in bold and the equivalent residues of RelP in regular font. The G-loop that stabilises the guanosine group of GTP/GDP and the product (p)ppGpp is labeled as well as the amphipathic α-helix α13 that contains a positive surface that coordinates the pyrophosphate group that is transferred from ATP to GDP or GTP. The comparison shows a remarkably conserved active despite both domains having less than 20% sequence identity. (**b**) Same as in (**a**) but displaying the ligands observed bound in the active site of both complexes; ppGpNp (in red) and AMP (in dark blue) in the case of the post-catalytic state complex of Rel$_{Tt}^{NTD}$ and GTP (in dark pink) and AMP (in light blue) in the case of the pre-catalytic state complex of RelP. From the structural comparison it becomes apparent that both enzymes most accommodate the two substrates in the same way with AMP and APCPP bound in almost the same position and orientation and ppGpNp accommodated with a small rearrangement of the G-loop likely to compensate for the lack of the extra phosphate group present in the GTP molecule. The catalytic residues D272, E345 and Q347 suggested to be involved in the hydrolysis and transferring of the pyrophosphate group are labeled together with the G-loop, α13 and the $^{\alpha}$P group of AMP.

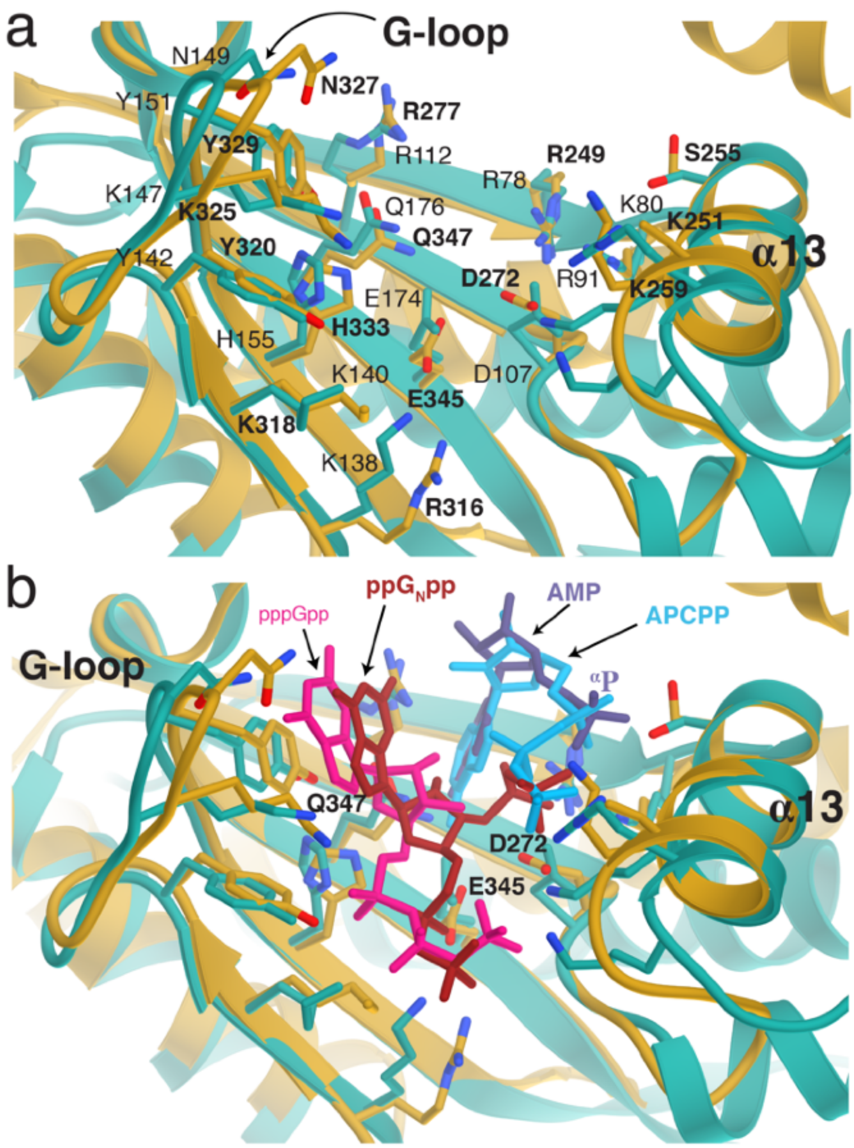

**Supplementary Fig. 8.** Single molecule FRET analysis. Two-dimensional (2D) histograms of the FRET efficiency E versus the stoichiometry S and PDA analysis of Rel$_{Tt}^{\mathrm{NTD}}{}_{6/287}$ in the presence of ppGpp (**a**), GDP+APCPP (**b**), APCPP (**c**), GDP (**d**) and GDP+ATP (**e**). Two-dimensional (2D) histograms of the FRET efficiency E versus the stoichiometry S and PDA analysis of Rel$_{Tt}^{\mathrm{NTD}}{}_{6/124}$ in the presence of ppGpp (**f**), APCPP (**g**) and ppGpp+EDTA (**h**).

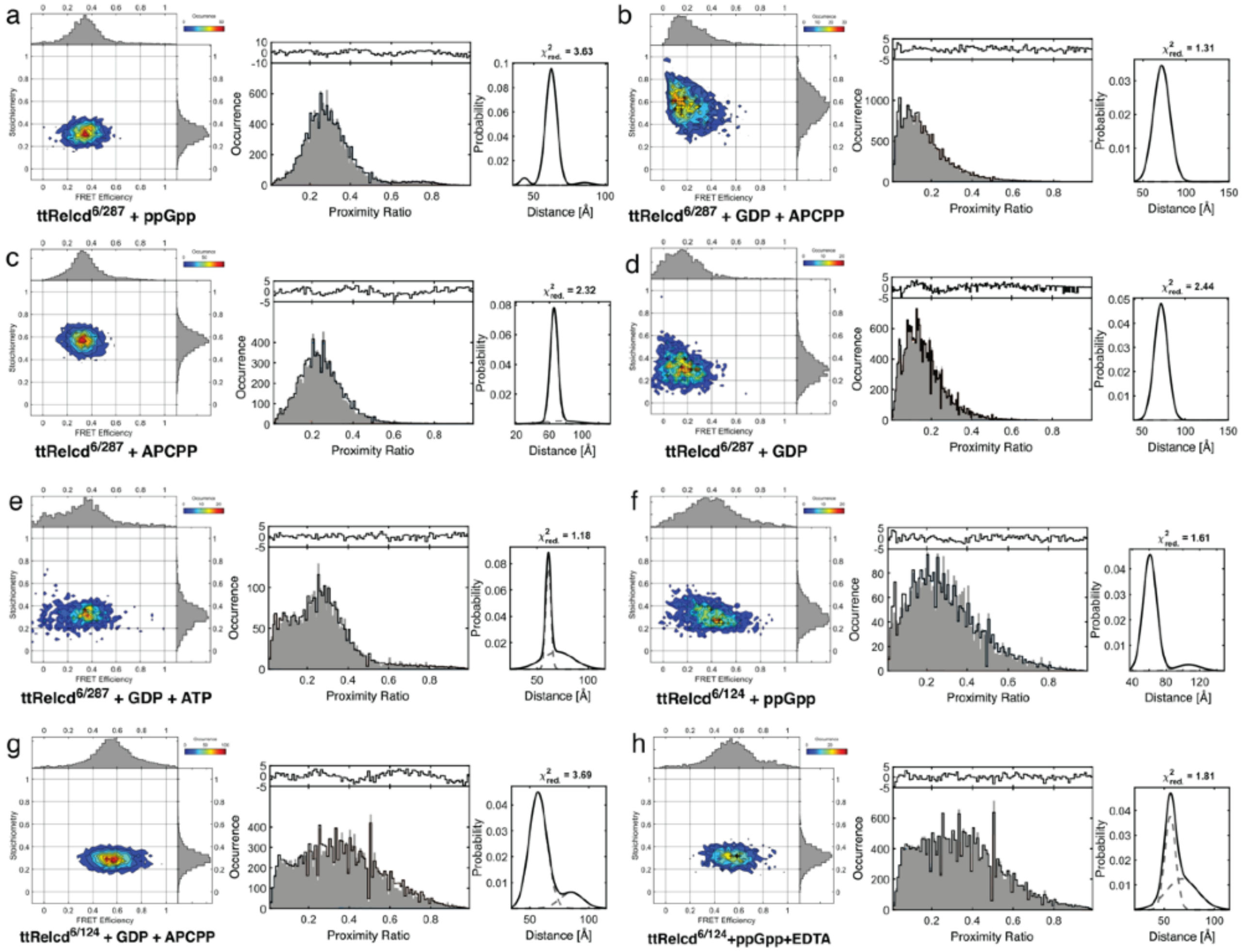

**Supplementary Fig. 9.** ITC measurements. (**a**) Titration of APCPP into $Rel_{Tt}^{NTD}$. (**b**) Theoretical thermodynamic parameters involved in the binding of APCPP to $Rel_{Tt}^{NTD}$ obtained from the differences between the binding of APCPP to $Rel_{Tt}^{NTD}$-GDP and the interaction of $Rel_{Tt}^{NTD}$ with GDP. All the thermodynamic parameters associated with each titration are listed in **Supplementary Table 2**.

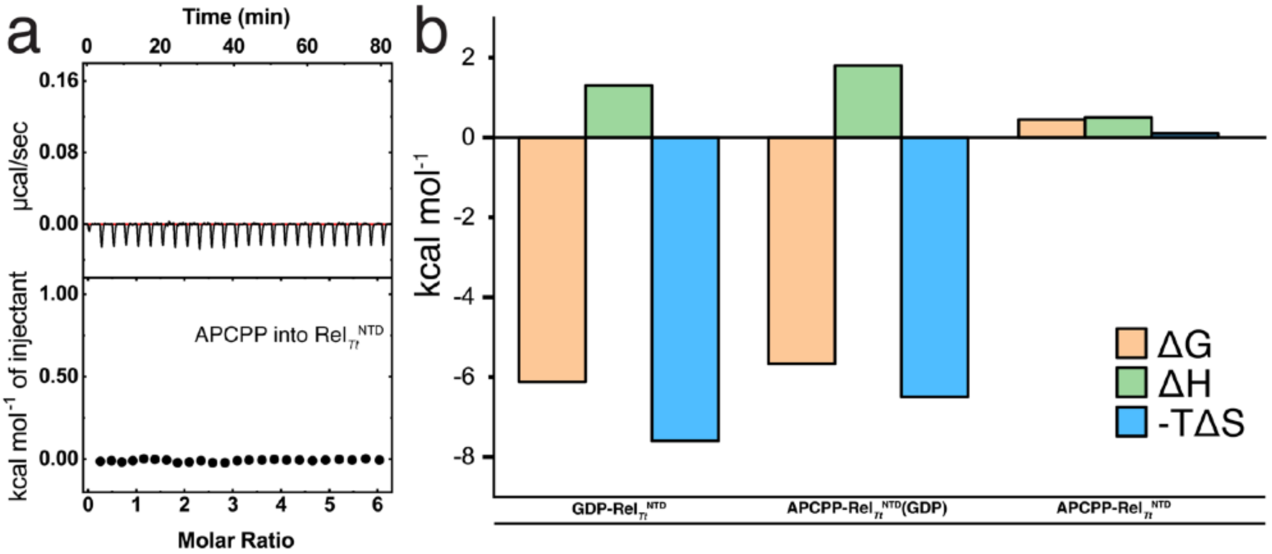
