## Supplementary material for "Nucleotide-mediated allosteric regulation of bifunctional Rel enzymes": Material and Methods

*Plasmids construction*

The chemically synthesized gene of the Rel*_Tt_* protein from *Thermus thermophilus* (GenScript) was cleaved with *XbaI* and *XhoI* restriction enzymes (NEB) and transferred to pET21b (Invitrogene), which was cut with the same restriction enzymes and phosphorylated with alkaline phosphatase (NEB). The sequence of the resulting plasmid, pET21b-HisTEV-ttRel, was verified by sequencing (Eurofins Genomics). To produce a plasmid for the overexpression of the catalytic (N-terminal) domains of the Rel*_Tt_* protein (Rel*_Tt_*^NTD^), the C-terminal part of the gene (from the residue V356) was removed by amplifying the entire plasmid with primers (Sigma Aldrich) flanking the sequence to be deleted (Supplementary table Y) with high fidelity Q5 polymerase (Sigma Aldrich). The acquired PCR reaction mixtures were treated with DpnI (NEB) to remove the template plasmid. Finally, the PCR products were purified with a PCR purification column (Sigma), phosphorylated with PNK (Sigma) and ligated. The ligation mixture was transferred to *E. coli* MC1061 by electroporation. The resulting plasmid pET21b-HisTEV-ttRelcd (residues 1-355) was sequenced (Eurofins Genomics).

The point mutations were introduced into the *rel_Tt_ (rel_Tt_^NTD^)* gene in pET21b-HisTEV-ttRel by amplifying the entire plasmid with high fidelity Q5 polymerase (Sigma Aldrich) using primer pairs (Sigma Aldrich) where one primer carries the desired mutation in its 5’end (**Supplementary Table 3**). The acquired PCR reactions mixtures were treated the same way as when constructing the C-terminal deletion of the enzyme. The removal and insertion of cysteine residues was carried out stepwise in the same way as described above for single mutations, using the plasmid acquired in the previous step as the template for the next one.

To facilitate the cloning of the hydrolysis active site mutants, the ligation mixtures of Rel*_Tt_*(R43A), Rel*_Tt_*(D81N/E), Rel*_Tt_*(R147G), Rel*_Tt_*(Y49F) and Rel*_Tt_*(N150A) were transformed to *E. coli* MC1061 already harbouring the plasmid pBAD33-Mesh1. To obtain this plasmid, the chemically synthesized MESH1 gene of *Homo sapiens* (GenScript) was amplified with high fidelity Q5 polymerase using primers Sac-RBS-Mesh-Fw and Mesh-Xho-Hind-Rev (**Supplementary Table 3**). The PCR product was cleaved using restriction enzymes HindIII and SacI (NEB) and transferred to the plasmid pBAD33 that was previously cleaved with the same enzymes and phosphorylated with alkaline phosphatase (NEB). The ligation mixture was transformed to *E. coli* MC1061 and verified by sequencing (Eurofins Genomics).

*Protein expression and preparation for purification*

HisTEV-Rel*_Tt_*, its catalytic domain (HisTEV-Rel*_Tt_^NTD^*) or its point mutations (HisTEV-Rel*_Tt_**) were expressed from the plasmids pET21b-HisTEV-Rel*_Tt_*(*) or pET21b-HisTEV- Rel*_Tt_^NTD^*, respectively. The plasmids were transferred to *E. coli* BL21(DE3) by electroporation. The cells carrying the plasmid were grown to an OD_600_ of 0.6 at 37 °C in LB supplemented with ampicillin (100 mg/l), prior to induction of protein expression by addition of IPTG (final concentration 0.5 mM) at 28 °C. The expression lasted for 5 h for Rel*_Tt_^NTD^*, or overnight for Rel*_Tt_*.

For the expression of the HisTEV-Rel*_Tt_** with point-mutations in the hydrolysis active centres (R43, Y49, D81, D146, R147 and N150) the pET21b-HisTEV-Rel*_Tt_** plasmids were transferred to *E. coli* BL21(DE3) cells that were already harbouring the plasmid pBAD33-Mesh1. The proteins were expressed as previously described, but besides ampicillin the growth media was supplemented also with chloramphenicol (10 mg/l) and L-arabinose (0.1%) to keep the Mesh1 expressed throughout the expression of the HisTEV-Rel*_Tt_^NTD^* hydrolysis mutant proteins. Next, the cells were collected and resuspended in resuspension buffer (50 mM Tris pH 8, 1.5 M KCl, 2 mM MgCl_2_, 1 mM TCEP) supplemented with cOmplete protease inhibitor cocktail (Roche). If not used immediately for downstream applications, the cells were flash-frozen in liquid nitrogen and stored at -80 °C. In every case cell disruption was performed using a cell cracker equilibrated with lysis buffer (50 mM Tris pH 8, 500 mM NaCl, 500 mM KCl, 1 mM TCEP) supplemented with complete protease inhibitor cocktail (Roche). The pellet was separated from the supernatant by centrifugation for 30 minutes at 30 000g.

The supernatant was loaded onto a gravity flow TALON-column, previously equilibrated with buffer A (25 mM Tris pH 8, 500 mM NaCl, 500 mM KCl, 10 mM MgCl_2_, 0.002% mellitic acid). The column was washed with 7 column volumes of buffer A and the bound protein eluted stepwise with buffers B1-3 (Buffer A containing 50 mM, 125 mM and 500 mM imidazole, respectively). The elution fractions containing His-Rel*_Tt_*^NTD^ were immediately concentrated using spin filters (Amicon) and loaded onto a GE Healthcare Superdex 200 16/60 gel filtration column equilibrated with GF buffer (50 mM Hepes pH 7.5, 500 mM NaCl, 500 mM KCl, 10 mM MgCl_2_, 0.002% mellitic acid; prior to crystallization procedures Tris pH 8 was used as a buffering agent). The protein fractions were checked for purity with SDS-PAGE. Pure protein was used immediately for further applications or stored in 35% glycerol at -20 °C for up to one week.

*Crystallization*

Before crystallization, the His-tag of all protein variants was cleaved off by adding TEV protease in a 1:200 molar ratio to the purified protein and incubating the mixture overnight at room temperature. The protein was analysed on SDS gel for purity and cleavage. Additionally, the removal of the tag was confirmed by Western Blotting using an anti-polyHis antibody (Sigma Aldrich). To clear Rel*_Tt_*^NTD^ from the His-tagged protease and free polyHis tag, the mixture was passed through a TALON column. The cleaved protein was subsequently concentrated for crystallization to 10-12 mg/ml. The concentration of the protein solution was estimated using the theoretical extinction coefficient at 280 nm of the proteins as implemented in the ProtParam tool.

Crystallization conditions were screened at 20 °C and 4 °C by the sitting-drop vapour-diffusion method. The drops were set up in Swiss (MRC) 96-well 2-drop UVP sitting-drop plates using the Mosquito HTS system (TTP Labtech). The drops consisting of 0.1 μl protein and 0.1 μl precipitant solution were equilibrated to 80 μl precipitant solution in the reservoir. Crystallization conditions were tested with several commercially available screens: Crystal Screen I and II (Hampton research), Helix, ProPlex, Pact Premier, JCSG, LMB and Morpheus II (Molecular Dimensions). For co-crystallization conditions of Rel*_Tt_*^NTD^ with nucleotides, Rel*_Tt_*^NTD^ at a concentration of 10-12 mg/ml was mixed with 100 mM of the nucleotides and incubated 10 minutes at room temperature. In every case the nucleotides were also added to the suitable cryo-protecting solution used for harvesting the crystals and the samples were vitrified in liquid N_2_ for storage and transport prior to X-ray exposure in the synchrotron.

X-ray diffraction data was collected at the SOLEIL synchrotron (Gif-sur-Yvette, Paris, France) on the Proxima 1 (PX1) and Proxima 2A (PX2A) beamlines using a PILATUS 6 M detector and an Eiger detector respectively. Because of the high anisotropic nature of the data from all the crystals we performed anisotropic cutoff and correction of the merged intensity data as implemented on the STARANISO server (http://staraniso.globalphasing.org/) using the DEBYE and STARANISO programs. In the case of the crystals of the Rel*_Tt_*^NTD^-ppGpp-AMP complex, the analysis of the data suggested a resolution of 2.95 Å (with 2.5 Å in a*, 3.0 in b* and 4.5 in c*). In the case of of the Rel*_Tt_*^NTD^-ppGp_N_p complex, the analysis of the data suggested a resolution of 2.75 Å (with 3.5 Å in a*, 3.5 in b* and 2.5 in c*).

*Structure determination*

The data were processed with the XDS suite^20^ and scaled with XSCALE or Aimless. In all cases, the unit-cell content was estimated with the program MATTHEW COEF from the CCP4 program suite^21^. Molecular replacement was performed with Phaser^22^. The crystals of the Rel*_Tt_*^NTD^-ppGpp complex diffracted on average to ~2.8 Å. We used the coordinates of Rel*_Seq_*^NTD^ as search model (PDBID 1VJ7)^23^. Because of the intrinsic interdomain dynamics of Rel catalytic domains we search for an MR solution using each catalytic domain as an independent ensemble. The MR solution from Phaser was used in combination with combined with Rosetta as implemented in the MR-Rosetta^24^ suit from the Phenix package^25^. MR-Rosetta could trace entirely the HD-domain and 70% of the SYN-domain. After several iterations of manual building with Coot^26^ and maximum likelihood refinement as implemented in Buster/TNT^27^, the model was extended to cover all the residues (R/Rfree of 17.9/22.9%).

In the case of the Rel*_Tt_*^NTD^-ppGpp-AMP complex, we used the coordinates of the individual HD and SYN domains from the structure of Rel*_Tt_*^NTD^-ppGpp as search model for MR in Phaser. The solution contained one molecule in the asymmetric unit. As with the Rel*_Tt_*^NTD^-ppGpp complex, we used MR-Rosetta^24^ after molecular replacement with Phaser. MR-Rosetta built the structure almost to completion and in the map resulting from MR-Rosetta, a clear density suggesting the presence of bound nucleotides in the SYN-domain was observed. The structure was completed after several iterations of manual building with Coot^26^ and maximum likelihood refinement as implemented in Buster/ Buster/TNT^27^ to an R/Rfree of 20.9/24.4%.

For the structure of Rel*_Tt_*^NTD^, we also used the coordinates of the individual HD and SYN domains from the structure of Rel*_Tt_*^NTD^-ppGpp as search model for MR in Phaser. In this case, the analysis of the unit cell content suggested the presence of 3 molecules in the asymmetric unit which was confirmed after MR. We used MR-Rosetta^24^ after molecular replacement with Phaser for automated model reconstruction. The structure was completed after several iterations of manual building with Coot^26^ and maximum likelihood refinement as implemented in Buster/ Buster/TNT^27^ to an R/Rfree of XX/XX%.

In every case the geometrical restraints of all small molecules were generated with the Grade Web Server (<http://grade.globalphasing.org>). **Supplementary Table X** details all the X-ray data collection and refinement statistics.

*Isothermal titration calorimetry*

All titrations were performed with an affinity ITC machine (TA instruments) at 15 °C. Stock solutions of APCPP (Jena biosciences) and GDP (Sigma Aldrich) of 650-670 mM were diluted in the protein buffer (50 mM Hepes pH 7.5; 500 mM KCl; 500 mM; NaCl; 10 mM MgCl_2_; 1 mM TCEP; 0.002 % mellitic acid) to a final concentration 1.6-2.4 mM. The purified Rel*_Tt_*^NTD^ was concentrated by ultrafiltration (Amicon ultra, 0,5 ml 30 kDa, Merck Millipore) to 80-100 μM. All final concentrations were verified by the absorption using a Nanodrop One (Thermo Scientific). All ITC measurements were performed by titrating 2 µl of the nucleotide into the protein using a constant stirring rate of 75 rpm. All data were processed and analysed using the NanoAnalyse and Origin software packages. **Supplementary Fig. 10** shows the representative results for each titration and the thermodynamic parameters derived from the analysis of the ITC data are shown in **Table 2**.

*Sample preparation for smFRET*

The Rel*_Tt_*^NTD^ variants (Rel*_Tt_*^NTD^_6/287_ and Rel*_Tt_*^NTD^_6/124_) suitable for smFRET measurements were prepared in 50 mM Hepes pH 7.5, 500 mM KCl; 500 mM NaCl; 10 mM MgCl2; 0.002 % mellitic acid. The Alexa Fluor 647 (Thermo Fisher Scientific) and Atto 488 (ATTO-TEC GmbH) used as acceptor and donor were covalently attached to the protein via Cys coupling. The concentrations of protein and dyes were determined by using a Nanodrop One (Thermo Scientific). For the labelling procedure the protein concentrated at 20-50 µM was incubated in a 50 µl reaction for 3 hours at room temperature (or overnight at 4 °C) with a 1.2-fold molar excess of each dye. After the labelling reaction, excess dye was remove with a Sephadex G-25 PD10 desalting column (GE healthcare), which was equilibrated with the buffer (50 mM HEPES pH 7.5; 500 mM KCl; 500 mM; NaCl; 10 mM MgCl_2_; 1 mM TCEP; 0.002 % mellitic acid). Labelled protein was concentrated (4-15 µM) with ultrafiltration (Amicon ultra, 0,5 ml 30 kDa, Merck Millipore) and stored at -20 °C in 20% glycerol.

The stored protein was always diluted a 1000-fold into the buffer prior to mixing with nucleotides in a reaction volume of 50 µl containing labelled protein (at a final concentration of 300 pM). The BSA-coated (1 mg/ml BSA) coverslip (Nunc Lab-Tek Chambered Coverglass, Thermo Fisher Scientific) was rinsed three times with the protein buffer (30 µl), prior to depositing a 30 µl drop of the prepared protein-nucleotide mixture. The background (needed for calculating E and S parameters, and for lifetime and PDA analysis) or scatter profile (needed for lifetime analysis) reference consisted of the same sample but without the protein or nucleotide. The small extra contribution of the nucleotides had a negligible effect. Measurements were performed at 22°C.

*smFRET data recording*

smFRET data were recorded on a homebuilt multiparameter fluorescence detection microscope with pulsed interleaved excitation (MFD-PIE) as established^28^, with minor modifications. Emission from a pulsed 483-nm laser diode (LDH-P-C-470, PicoQuant) was cleaned up (Chroma ET485/20x, F49-482; AHF analysentechnik AG), emission from a 635-nm laser diode (LDH-P-C-635B, PicoQuant) was cleaned up (Chroma z635/10x, PicoQuant), and both lasers were alternated at 26.67 MHz (PDL 828 Sepia II, PicoQuant), delayed ~18 ns with respect to each other, and combined with a 483-nm re- flecting dichroic mirror in a single-mode optical fiber (coupler, 60FC-4- RGBV11-47; fiber, PMC-400Si-2.6-NA012-3-APC-150-P, Schäfter + Kirchhoff GmbH). After collimation (60FC-L-4-RGBV11-47, SuK GmbH), the linear polarization was cleaned up (CODIXX VIS-600- BC-W01, F22-601; AHF analysentechnik AG), and the light (100 mW of 483-nm light and 50 mW of 635-nm light) was reflected into the back port of the microscope (IX70, Olympus Belgium NV) and upward [3-mm-thick full-reflective Ag mirror, F21-005 (AHF) mounted in a total internal reflection fluorescence filter cube for BX2/IX2, F91-960; AHF analysentechnik AG] to the objective (UPLSAPO-60XW, Olympus). Sample emission was transmitted through a 3-mm-thick excitation polychroic mirror (Chroma zt470-488/640rpc, F58-PQ08; AHF anal- ysentechnik AG), focused through a 75-mm pinhole (P75S, Thorlabs) with an achromatic lens (AC254-200-A-ML, Thorlabs), collimated again (AC254-50-A-ML, Thorlabs), and spectrally split (Chroma T560lpxr, F48-559; AHF analysentechnik AG). The blue range was filtered (Chroma ET525/50m, F47-525, AHF analysentechnik AG), and polarization was split (PBS251, Thorlabs). The red range was also filtered (Chroma ET705/100m, AHF analysentechnik AG), and polarization was split (PBS252, Thorlabs). Photons were detected on four avalanche photo- diodes (PerkinElmer or EG&G SPCM-AQR12/14), which were connected to a time-correlated single-photon counting (TCSPC) device (SPC-630, Becker & Hickl GmbH) over a router (HRT-82, Becker & Hickl) and power supply (DSN 102, PicoQuant). Signals were stored in 12-bit first-in-first-out (FIFO) files. Microscope alignment was carried out using fluorescence correlation spectroscopy (FCS) on freely diffusing ATTO 488-CA and ATTO 655-CA (ATTO-TEC) and by connecting the detectors to a hardware correlator (ALV-5000/EPP) over a power splitter (PSM50/51, PicoQuant) for alignment by real-time FCS. Data were loaded in the PAM software (D. C. Lamb, Ludwig-Maximilians-Universität Munich) written in MATLAB (MathWorks). Instrument response functions (IRFs) were recorded in a solution of ATTO 488-CA or ATTO 655-CA in near-saturated centrifuged potassium iodide at a 25-kHz average count rate. Macrotime-dependent microtime shifting was present and corrected for two (blue/parallel and red/perpendicular) of four avalanche photodiodes (APDs) on the instrument response function (IRF) data. Signals from each TCSPC channel were divided in time gates to discern 483-nm excited FRET photons from 635-nm excited acceptor photons. A two-color MFD all-photon burst search algorithm using a 500-ms sliding time window (minimum of 50 photons per burst, minimum of 5 photons per time window) and a kernel density estimator (ALEX-2CDE < 12) were used to identify single donor- acceptor–labeled molecules in the fluorescence trace.

*Photon distribution analysis (PDA)*

Static PDA was carried out to obtain the absolute interdye distance distribution as described before, assuming two (or more) Gaussian distributed states^29^. Practically, for each FRET data set, raw bursts were re-binned in different time bins (0.2, 0.5, 0.75, and 1 ms), and four histograms were constructed and analyzed simultaneously. Data were plotted in a FRET efficiency versus stoichiometry plot to deselect bins with complex acceptor photophysics, and only bins with at least 20 and maximally 250 photons (to reduce calculation time) were used for PDA analysis. A three-state model for a Gaussian distance distribution was used to generate a library of simulated EPR values, which was subsequently fitted to the experimental *E*_PR_ histogram using a reduced χ^2^–guided simplex search algorithm. The mean and width of all Gaussian distributed sub-states were globally optimized, whereas the state area A was globally optimized over a single sample. Moreover, the standard deviation of the distance distributions was globally optimized at a fraction *F* of the corresponding distance to increase fitting robustness, which has been shown before to be reasonable for FRET experiments with a blinking FRET acceptor. Finally, *F* was globally optimized over all states and data sets^30^. A probability density function (PDF) was calculated per state using the R and s parameters obtained from PDA analysis that describe the underlying Gaussian distributed states. The summed PDF was scaled to a total area of unity, with the PDF area of each state scaled to the corresponding fraction of molecules. Criteria for a good fit were a low reduced χ^2^ value, as well as a weighted residuals plot free of trends.

*Preparation of* T. thermophilus *70S ribosomes*

70S ribosomes were prepared from *T. thermophilus* strain HB8 as described^31^ with minor modifications. Approximately 20 g of frozen *T. thermophilus* cell paste was resuspended in 50 ml of buffer A (100 mM NH_4_Cl, 10.5 mM Mg-acetate, 0.5 mM EDTA, 6 mM β-mercaptoethanol, 20 mM Tris:HCl pH 7.5, NB! β-mercaptoethanol should be added directly prior to use) supplemented with 1 mU Turbo DNase (Thermo Fisher Scientific), 0.1 mM PMSF, and opened by three passages on a high-pressure cell disrupter (Stansted Fluid Power) at 220 MPa. Lysed cells were clarified by centrifugation for 60 min at 30,000 rpm (Ti 45 rotor, Beckman). The supernatant was carefully removed, taking care not to disturb the pellet of cell debris and then carefully applied on sucrose cushion (1.1 M sucrose, 0.5 M NH_4_Cl, 15 mM Mg-acetate, 0.5 mM EDTA, 3 mM β-mercaptoethanol, 20 mM Tris:HCl pH 7.5) by loading 15 ml of supernatant on the top of 40 ml of sucrose cushion in 45Ti centrifuge tubes. After centrifugation for 8 hours at 35,000 rpm (Ti 45 rotor, Beckman) the ribosomes were pelleted and the supernatant was removed. This is a crucial step: light jelly-like debris should be carefully removed by tipping the tubes over, yielding clear ribosome pellets. If necessary, the pellets can be washed with ice-cold buffer A. Ribosomal pellets were resuspended in the buffer A and then applied to the sucrose cushions (45Ti centrifuge tubes), and centrifuged for 18 hours at 28,000 rpm (Ti 45 rotor, Beckman). Resulting pellets were resuspended in the buffer A and resolved on a 10-40% sucrose gradient in overlay buffer (60 mM NH_4_Cl, 15 mM Mg(OAc)_2_, 0.25 mM EDTA, 3 mM β-mercaptoethanol, 20 mM Tris:HCl pH 7.5) in a zonal rotor Ti 15 (Beckman) (17 hours at 21,000 rpm). The peak containing pure 70S ribosomes was pelleted by centrifugation (20 hours at 35,000 rpm), and the final ribosomal preparation was dissolved in HEPES:Polymix buffer^32^ (20 mM HEPES:KOH pH 7.5, 2 mM DTT, 5 mM Mg(OAc)_2_, 95 mM KCl, 5 mM NH_4_Cl, 0.5 mM CaCl_2_, 8 mM putrescine, 1 mM spermidine. 70S concentration was measured spectrophotometrically (1 OD_260_ corresponds to 23 nM of 70S) and ribosomes were aliquoted (50-100 μl per aliquot), snap-frozen in liquid nitrogen and stored at –80°C.

*Preparation of* T. thermophilus *70S initiation complexes (70S IC)*

Initiation complexes were prepared by as established^33^, with minor modifications. The reaction mix containing *T. thermophilus* 70S ribosomes (final concentration of 1 μM) with 1 μM *E. coli* IF1, 1 μM IF2, 1 μM IF3, 1.5 μM ^3^H-fMet-tRNA_i_^fMet^, 1.5 μM mRNA MVFStop (5′-GGC**AAGGAGGA**GAUAAGAAUGGUUUUCUAAUA-3′), 1 mM GTP and 2 mM DTT in HEPES:Polymix buffer with 5 mM Mg(OAc)_2_ was incubated at 37 °C for 30 min. Then the ribosomes were pelleted through a sucrose cushion (1.1 M sucrose in HEPES:Polymix buffer with 15 mM Mg^2+^) at 50,000 rpm for 2 hours (TLS-55, Beckman), the pellet was dissolved in HEPES:Polymix buffer (5 mM Mg(OAc)_2_), aliquoted, snap-frozen in liquid nitrogen and stored at –80 °C.

*Biochemical assays*

*H^3^-ppGpp synthesis assays*: experiments were performed as described for *E. coli*^34^ with minor modifications. The reaction mixtures typically contained 120 nM *T. thermophilus* 70S IC(MV), 30 nM Rel, guanosine nucleoside substrate (300 μM H^3^-GDP), (ParkinElmer) *E. coli* tRNA^Val^, all in HEPES:Polymix buffer at 5 mM Mg^2+^ final concentration. After preincubation at 40°C for 3 minutes, the reaction was started by the addition of prewarmed ATP to the final concentration of 1 mM, and 5 μl aliquots were taken throughout the time course of the reaction and quenched with 4 μl 70% formic acid supplemented with a cold nucleotide standard (4 mM GDP and 4 mM GTP) for UV-shadowing. Individual quenched timepoints were spotted PEI-TLC plates (MACHEREY-NAGEL) and nucleotides were resolved in 0.5 KH_2_PO_4_ pH 3.5 buffer. The TLC plates were dried, cut into sections as guided by UV-shadowing, and ^3^H radioactivity was quantified by scintillation counting in an Optisafe-3 (Fisher) scintillation cocktail.

*H^3^-ppGpp hydrolysis assays*: the reaction mixtures typically contained 100 nM Rel*_Tt_*, guanosine nucleoside substrate (500 μM H^3^-ppGpp), 0.5 mM MnCl_2_, all in HEPES:Polymix buffer at 5 mM Mg^2+^ final concentration. After preincubation at 40 °C for 3 minutes, the reaction was started by the addition of prewarmed Rel and 5 μl aliquots were taken throughout the time course of the reaction and quenched with 4 μl 70% formic acid supplemented with a cold nucleotide standard (4 mM GDP) for UV-shadowing.

**Online References:**

20.	Kabsch, W. Xds. *Acta Crystallogr D Biol Crystallogr* **66**, 125-32 (2010).

21.	Collaborative Computational Project, N. The CCP4 suite: programs for protein crystallography. *Acta Crystallogr D Biol Crystallogr* **50**, 760-3 (1994).

22.	McCoy, A.J. *et al.* Phaser crystallographic software. *J Appl Crystallogr* **40**, 658-674 (2007).

23.	Hogg, T., Mechold, U., Malke, H., Cashel, M. & Hilgenfeld, R. Conformational antagonism between opposing active sites in a bifunctional RelA/SpoT homolog modulates (p)ppGpp metabolism during the stringent response [corrected]. *Cell* **117**, 57-68 (2004).

24.	Terwilliger, T.C. *et al.* phenix.mr_rosetta: molecular replacement and model rebuilding with Phenix and Rosetta. *J Struct Funct Genomics* **13**, 81-90 (2012).

25.	Afonine, P.V. *et al.* Towards automated crystallographic structure refinement with phenix.refine. *Acta Crystallogr D Biol Crystallogr* **68**, 352-67 (2012).

26.	Emsley, P. & Cowtan, K. Coot: model-building tools for molecular graphics. *Acta Crystallogr D Biol Crystallogr* **60**, 2126-32 (2004).

27.	Smart, O.S. *et al.* Exploiting structure similarity in refinement: automated NCS and target-structure restraints in BUSTER. *Acta Crystallogr D Biol Crystallogr* **68**, 368-80 (2012).

28.	Talavera, A. *et al.* Phosphorylation decelerates conformational dynamics in bacterial translation elongation factors. *Sci Adv* **4**, eaap9714 (2018).

29.	Antonik, M., Felekyan, S., Gaiduk, A. & Seidel, C.A. Separating structural heterogeneities from stochastic variations in fluorescence resonance energy transfer distributions via photon distribution analysis. *J Phys Chem B* **110**, 6970-8 (2006).

30.	Kalinin, S. *et al.* A toolkit and benchmark study for FRET-restrained high-precision structural modeling. *Nat Methods* **9**, 1218-25 (2012).

31.	Polikanov, Y.S., Steitz, T.A. & Innis, C.A. A proton wire to couple aminoacyl-tRNA accommodation and peptide-bond formation on the ribosome. *Nat Struct Mol Biol* **21**, 787-93 (2014).

32.	Antoun, A., Pavlov, M.Y., Tenson, T. & Ehrenberg, M.M. Ribosome formation from subunits studied by stopped-flow and Rayleigh light scattering. *Biol Proced Online* **6**, 35-54 (2004).

33.	Murina, V., Kasari, M., Hauryliuk, V. & Atkinson, G.C. Antibiotic resistance ABCF proteins reset the peptidyl transferase centre of the ribosome to counter translational arrest. *Nucleic Acids Res* **46**, 3753-3763 (2018).

34.	Kudrin, P. *et al.* The ribosomal A-site finger is crucial for binding and activation of the stringent factor RelA. *Nucleic Acids Res* **46**, 1973-1983 (2018).
