## Supplementary Tables for "Nucleotide-mediated allosteric regulation of bifunctional Rel enzymes"

**Supplementary Table 1. ppGpp synthesis and hydrolysis reaction parameters. Each measurement was performed in triplicates.**

|  | **Synthase activity** | | | | | | **Hydrolase activity** |
| --- | --- | --- | --- | --- | --- | --- | --- |
|  | **Substrate : GDP+ATP** | | | **Substrate : GDP+APPNP** | | | **Substrate : ppGpp** |
|  | **Rel*_Tt_*** | **Rel*_Tt_*+IC** | **Rel*_Tt_*+IC+tRNA^Val^** | **Rel*_Tt_*** | **Rel*_Tt_*+IC** | **Rel*_Tt_*+IC+tRNA^Val^** | **Rel*_Tt_*** |
| Rel*_Tt_* | 4.2±0.7 | 37.7±1.9 | 332.3±41.7 | N.D. | N.D. | 6.7±2.4 | 6.3±1.0 |
| R249AD272A | N.D. | 8.5±1.0 | 12.0±2.0 | - | - | - | - |
| R249AR277A | 2.6±0.6 | 8.6±2 | 10.5±1.8 | - | - | - | - |
| R249AD272AY329A | N.D. | 10.9±3.6 | 8.8±0.69 | - | - | - | - |
| R249AR277AY329A | N.D. | 4.5±2.3 | 5.4±2.1 | - | - | - | - |
| R43A | - | - | - | - | - | - | N.D. |
| Y49F | - | - | - | - | - | - | N.D. |
| E80Q | - | - | - | - | - | - | N.D. |
| D81N | - | - | - | - | - | - | N.D. |
| D81E | - | - | - | - | - | - | N.D. |
| R147G | - | - | - | - | - | - | N.D. |
| N150A | - | - | - | - | - | - | N.D. |

**Supplementary Table 2. Binding parameters obtained from the ITC titrations.** The binding affinities were determined from fitting a single interaction model to the experimental ITC data according to the experimental setup. Data represent mean values ± s.d.

| **Titration** | **Kd (μM)** | **ΔG (kcal mol^-1^)** | **ΔH (kcal mol^-1^)** | **-TΔS (kcal mol^-1^)** | **n** |
| --- | --- | --- | --- | --- | --- |
| GDP into Rel*_Tt_*^NTD^ | 23.0 ± 0.5 | -6.12 ± 0.01 | 1.3 ± 1 | -7.6 ± 1 | 3 |
| GDP into Rel*_Tt_*^NTD^-APCPP complex | 24.7 ± 0.9 | -6.1 ± 0.1 | 0.4 ± 1 | -6.5 ± 1 | 3 |
| APCPP into ttRelcd | - | - | - | - | 2 |
| APCPP into Rel*_Tt_*^NTD^-GDP complex | 50.4 | -5.67 ± 0.07 | 1.8 ± 1 | -7.5± 1 | 3 |

**Supplementary Table 3. Oligonucleotides used for the construction of recombinant Rel*_Tt_* and all the Rel*_Tt_* substitutions**

| **Oligo name** | **sequence** | **purpose** |
| --- | --- | --- |
| Rel-Tthe-CD | 5’-GCGATGCATTTCGCGGGTACGA-3’ | N-terminal deletion of Rel*_Tt_* |
| F-pET-stop | 5’-TGAGATCCGGCTGCTAACAAAGCCC-3’ |  |
| TT-L6-Rev | 5’-GTCCGCACCCACCATGCCC-3’ | L6C substitution of Rel*_Tt_* or Rel*_Tt_*^NTD^ |
| TT-L6C-Fw | 5’-**TGC**GGTCTGTGGAACCGCCTGGAAC-3’ |  |
| TT-C82-Fw | 5’-GAAGAACTGGAACGTCGCTTTGGTC-3’ | C82S substitution of Rel*_Tt_* or Rel*_Tt_*^NTD^ |
| TT-C82S-Rev | 5’-CGGCGCA**ACG**CCGCTGTCTTCCAGCGTATCATGCAGCAG-3’ |  |
| TT-E124-Fw | 5’-GATCTGCGCCAGATGTTCATTGC-3’ | E124C substitution of Rel*_Tt_* or Rel*_Tt_*^NTD^ |
| TT-E124C-Rev | 5’- **GCA**GGCGCGACGTTCTTCACC-3’ |  |
| TT-T287-Rev | 5’-CGGTGCCGGTTTCGGGTC-3’ | T287C substitution of Rel*_Tt_* or Rel*_Tt_*^NTD^ |
| TT-T287C-Fw | 5’- **TGC**CGCGAATCGCAAGCTCTGCG-3’ |  |
| TT-C299-Fw2 | 5’-GGGTCTGGTTCACGCACTGTG-3’ | C299A substitution of Rel*_Tt_* or Rel*_Tt_*^NTD^ |
| TT-C299A-Rev | 5’- AGCACATGATA**CGC**GACCTGTTTTTCACGCAGAGC-3’ |  |
| TT-R43A-Fw | 5’-**GCG**CGCTCTGGCGAACCGTATATCAC-3’ | R43A substitution of Rel*_Tt_* |
| TT-R43-Rev | 5’-CAGCTGACCACGATGAGCTTC-3’ |  |
| TT-Y49-Rev | 5’-CGGTTCGCCAGAGCGACG-3’ | Y49F substitution of Rel*_Tt_* |
| TT-Y49F-Fw | 5’-**TTT**ATCACCCACCCGGTCGCAG-3’ |  |
| TT-D81E-Rev | 5’-CGGCGCAACGCCGCA**TTC**TTCCAGCGTATCATGCAGCAG-3’ | D81E substitution of Rel*_Tt_*, with TT-C82-Fw |
| TT-D81N-Rev | 5’-CGGCGCAACGCCGCA**GTT**TTCCAGCGTATCATGCAGCAG-3’ | D81N substitution of Rel*_Tt_*, with TT-C82-Fw |
| TT-R147G-Fw | 5’-GAC**GGC**CTGCATAATCTGCGTACCC-3’ | R147G substitution of Rel*_Tt_* |
| TT-D146/7-Rev | 5’-TGCCAGTTTAACAATGATAATACGCAC-3’ |  |
| TT-N150A-Fw | 5’-**GCG**CTGCGTACCCTGGAACACATG-3’ | N150A substitution of Rel*_Tt_* |
| TT-N150-Rev | 5’-ATGCAGGCGGTCTGCCAG-3’ |  |
| TT-R249A-Fw | 5’-**GCC**CCGAAACACCTGTATTCAATTTGG-3’ | R249A substitution of Rel*_Tt_*, Rel*_Tt_*^D272A^ and Rel*_Tt_*^R277A^ |
| TT-R249-Rev | 5’-ACCCGTCACTTCAAAGCCC-3’ |  |
| TT-D272A-Fw | 5’-**GCT**CTGCTGGCGGTTCGCGTC-3’ | D272A substitution of Rel*_Tt_* |
| TT-D272-Rev | 5’-GTAGATCTGTTCCAGGGTTTTGCC-3’ |  |
| TT-R277-Fw | 5’-GTCATTCTGGACCCGAAACCGG-3’ | introducing the mutation R277A to Rel*_Tt_* |
| TT-R277A-Rev | 5’-**CGC**AACCGCCAGCAGATCGTAGATC-3’ |  |

**Supplementary Table 4. Data collection and processing.** The CC1/2 criterion was used to determine the resolution range. Values for the outer shell are given in parentheses.

| **Sample** | **Rel*_Th_*^NTD^** | **Rel*_Th_*^NTD^-ppGpp** | **Rel*_Th_*^NTD^-ppGp_N_p-AMP** |
| --- | --- | --- | --- |
| Diffraction source | Soleil PX1 | Soleil PX1 | Soleil PX2A |
| Wavelength (Å) | 1.008 | 1.008 | 0.9801 |
| Space group | P 4_1_ 2_1_ 2 | P 4_1_ 2 2 | C2 |
| *a*, *b*, *c* (Å) | 105.7 105.7 241.4 | 88.4 88.4 182.7 | 120.8 50.3 86.1 |
| α, β, γ (°) | 90.0 90.0 90.0 | 90.0 90.0 90.0 | 90.0 111.0 90.0 |
| Resolution range (Å) | 52.87 - 2.89 (2.998 - 2.89) | 51.60 - 2.75 (2.85 - 2.75) | 56.36 - 2.95 (3.06 - 2.95) |
| Total N^o^. of reflections | 297005 (1413) | 156684 (2496) | 67782 (2399) |
| N^o^. of unique reflections | 21371 (124) | 12385 (223) | 6572 (255) |
| Completeness after anisotropic correction (%) | 93.7 (90.9) | 94.7 (97.9) | 91.9 (94.8) |
| Redundancy | 13.9 (11.4) | 12.7 (11.2) | 10.3 (9.4) |
| 〈*I*/σ(*I*)〉 | 14.38 (1.31) | 10.15 (1.21) | 13.80 (2.23) |
| *CC*_1/2_ | 0.999 (0.724) | 0.996 (0.611) | 0.996 (0.868) |
| *R*_r.i.m._ | 0.12 (1.57) | 0.21 (1.69) | 0.12 (1.1) |
| *R*_pim_ | 0.03 (0.56) | 0.06 (0.52) | 0.04 (0.36) |
| R-factor (%) | 26.4 | 17.9 | 20.9 |
| R_free_-factor (%) | 31.1 | 23.0 | 24.4 |
| Ramachandran profile |  |  |  |
| Core | 93.2 | 96.6 | 93.8 |
| Allowed | 6.5 | 3.4 | 6.2 |
| Outliers | 0.3 | 0.0 | 0.0 |
| R.m.s. deviations |  |  |  |
| Bond lengths (Å) | 0.005 | 0.014 | 0.015 |
| Bond angles (°) | 0.76 | 1.83 | 1.90 |
| Number of atoms | 7537 | 2913 | 2681 |
| Macromolecules | 7531 | 2778 | 2617 |
| Solvent | 1 | 97 | 20 |
| Ligands | 5 | 38 | 44 |
| B-factors (Å2) |  |  |  |
| All atoms | 83.09 | 65.1 | 100.79 |
| Macromolecules | 83.07 | 64.1 | 100.73 |
| Solvent atoms | 123.59 | 179.5 | 45.04 |
| Ligands | 45.87 | 48.5 | 129.32 |
| PDB ID | XXXX | XXXX | XXXX |
